# Unfolded to Folded: Unraveling the Secrets of Protein Folding with ProteusFold

**DOI:** 10.1101/2025.10.08.679099

**Authors:** Saleh Sakib Ahmed

**Affiliations:** Computer Science and Technology, BUET, West Palashi, Dhaka, 1000, Dhaka, Bangladesh

**Keywords:** Protein Folding, Light-weight, Transformer, Protein Structure Tokenization, Folding hotspot

## Abstract

Protein folding has long been regarded as the “holy grail” of biology, typically demanding large models and massive GPU clusters. This study introduces ProteusFold, a compact and interpretable model with only 993,408 parameters that achieves state-of-the-art accuracy on modest hardware with an inference time of 0.0011 seconds. By framing folding as an unfolded-to-folded sequence transformation using a novel structural tokenization, ProteusFold reduces regression complexity while preserving bond connectivity through the concept of “Synapses.” It achieves near-atomic fidelity (RMSD 0.24,Å, GDT-TS 99.85) and excels in protein–protein docking with a mean DockQ of 0.7675, with 95.5% of complexes above the 0.23 threshold. Compared to AlphaFold2’s Predicted Aligned Error (6.28), ProteusFold attains 0.396, representing an order-of-magnitude gain in positional accuracy. Beyond accuracy and efficiency, the model provides residue-level attribution analyses that highlight biologically significant residues, serving as a preliminary guide for experiments. Furthermore, ProteusFold is the first to provide atomic-level attribution of key electronic and thermal properties, offering deeper insight into folding mechanisms and pinpointing the specific atoms responsible for distinct scenarios. Moreover, a meta-analysis suggests the presence of *folding hotspots*, where critical residues cluster, revealing new avenues for discovery. Thus, ProteusFold delivers accuracy, interpretability, and efficiency, broadening access to protein-folding research.

## 1 Introduction

The word *protein* derives from the Greek term “*protios*,” meaning “primary,” underscoring their essential role in nearly all biological processes. Proteins catalyze reactions as enzymes, provide structural support to cells and tissues, transport and store molecules, regulate signaling pathways, and defend the body through the immune system [1, 2]. Their central role in these processes makes them indispensable to life, yet they possess a unique property. Like the Greek god Proteus, who could change form at will, proteins fold from linear chains into diverse three-dimensional structures that define their function and behavior [3]. Predicting this folded structure directly from sequence is known as the “Protein Folding Problem.”

For decades, the protein folding problem baffled scientists and stood as one of the greatest challenges in biology and computer science. Although early efforts [4, 5] provided important insights, they failed to produce a general solution. This changed in 2021 with the release of AlphaFold2 [6], a 21M-parameter non-generative model [7] that achieved unprecedented accuracy. Implemented as a five-model ensemble, AlphaFold2 integrates MSAs, pairwise features, and structural templates, while applying axial and triangular attention to iteratively predict backbone coordinates^1^ through an eight-block invariant-point-attention module. Afterward, side chains are reconstructed using a template-based approach, and predictions are refined through three recycling iterations^2^, followed by 200 steps of AMBER relaxation [8] to optimize local geometry and remove stereochemical violations. Regarding computational requirements, training took 11 days on 128 TPUv3 devices [7] using 256-residue crops with fine-tuning on 384, whereas inference for a single protein typically takes about ten minutes [9]. Following this breakthrough, multiple large-scale models were developed, each requiring substantial GPU resources and prolonged training periods. For example, among MSA-based methods, RosettaFold [10] (130M parameters) was trained for 200 epochs on eight 32GB V100 GPUs over four weeks, relying on both MSAs and structural templates. By contrast, protein language model (PLM)^3^ approaches, such as ESM-Fold [11], leverage a 15-billion-parameter PLM to predict structures directly from single sequences, thereby eliminating the need for MSAs. The model was fully trained in approximately two weeks on a heterogeneous cluster of 2,000 GPUs [11]. Furthermore, to broaden accessibility of these massive models, open-source reimplementations like ColabFold [12] and OpenFold [13] were introduced, though high computational demands and slow inference still remained.

In 2024, DeepMind [14] introduced AlphaFold3 [15], a diffusion-based model that still relied on MSAs, templates, and pairwise features, but expanded capabilities to accurately predict protein–protein, protein–RNA, and other biomolecular complexes. This achievement contributed to two of its authors receiving the 2024 Nobel Prize. Since then, models have grown even larger across multiple paradigms: ultra-large PLMs such as XtrimopGLM [16] (100B parameters; 1 trillion tokens) and ESM3 (98B parameters); MSA-based RosettaFold All-Atom [17] (83M parameters); and generative models like Proteina [18] (400M parameters) are some mentionable ones. Meanwhile, efforts to accelerate these computationally demanding systems using supercomputers [19, 20] reinforced the perception that protein folding has been “solved”—but only with massive computational resources.

This paradigm presents two critical limitations. First, the computational cost of training and large-scale inference places these tools beyond the reach of many researchers. Second, reliance on MSAs and single sequence introduces several challenges. MSAs are often noisy and error-prone due to misalignments, may be unavailable for rapidly evolving or poorly studied protein families, and are computationally expensive to generate at scale [21]. Moreover, MSA quality depends heavily on the size and diversity of reference databases, introducing sampling bias and leaving proteins with sparse homologs poorly represented [22]. Protein language models face a similar issue, as they are biased by unequal sequence sampling driven by uneven species representation [23]. MSAs can also blur functional signals by averaging across divergent sequences, masking biologically relevant variations such as adaptive mutations or disorder-to-order transitions. Finally, constructing MSAs is itself a computational bottleneck, limiting their utility in high-throughput or real-time applications.

Beyond MSA reliance and heavy computational requirements, most large-scale models remain closed systems with limited interpretability, despite recent efforts such as ExplainFold [24]. Deep learning models have the potential to be a source of great guidance for researchers to find preliminary guiding points for experimentation by identifying crucial residues. However, none of these models explicitly incorporate the atomic-level physicochemical properties of the residues, making the interpretation of model difficult. At the same time, interpreting predictions involving regression of structures is challenging, as an increase or decrease in the value of the predicted structure cannot be meaningfully interpreted. In this study, these limitations were mitigated by eliminating the dependence on MSAs and redefining the protein folding problem for interpretable results.

### 1.1 Redefining Protein Folding

At first glance, a folded protein appears overwhelmingly complex, making the prediction of atomic coordinates from a linear residue sequence seem intractable. To address this, Proteus simulates this process as it occurs in nature: unfolded proteins—readily generated by existing tools [25]—are translated into their folded states using atomic properties as guiding information. The main challenge in this endeavor lies in the large conformational gap between unfolded and folded states, which, in principle, requires immense computational resources. At the same time, the model must ensure that the bond lengths and angles remain physically valid.^4^ ProteusFold addresses these challenges by decomposing the protein into standardized residue tokens, each with a “synapse” (details in Sec. 2.1) that anchors it to its neighbors by retaining the start coordinates of the subsequent residues. Each token, folded or unfolded, is then shifted to the origin to standardize the representation. Thus, in this formulation, the folding problem becomes the task of translating standardized residue tokens from the unfolded to the folded space, guided by atomic properties. Finally, the predictions are reassembled through these synapses to restore the global coordinates of each residue, thereby preserving the bond physics and connectivity of each residue. This is a simple, yet powerful strategy that preserves physical validity while greatly reducing computational cost and converting the folding problem into a tractable regression. This enabled the development of a model that can be trained in a matter of hours for less than one US dollar and under 1 kWh of energy, with large-scale inference taking only seconds and a single protein prediction requiring approximately 0.0011 seconds. In contrast, competing approaches typically require hundreds of thousands of dollars, several thousand kWh, and days of training and inference. An approximate cost analysis, quantifying both monetary and energy savings relative to existing models, is presented in Sec. S9 of the Supplementary.

### 1.2 Interpretation and Applicability

Unlike conventional interpretation methods that focus solely on model predictions, ProteusFold derives interpretability directly from the loss function. This approach reveals how individual features either decrease the loss—thereby moving closer to the reference structure—or increase it, causing deviation. Such a formulation enables both residue- and atomic-level interpretation of folding dynamics by quantifying their influence on the loss landscape.

By providing biologically grounded residue-level attributions, ProteusFold offers experimentally supported insights that can serve as preliminary guidance for future protein engineering and mutagenesis studies. Moreover, for the first time, atomic-level interpretations of fundamental electronic and thermal properties are made accessible within a folding framework, providing deeper mechanistic understanding and pinpointing the specific atoms responsible for distinct structural transitions.

To the best of our knowledge, ProteusFold is the first method to deliver atomic-scale interpretability in protein folding. Additionally, positional analysis of attribution patterns suggests the existence of “folding hotspots”—regions where critical residues consistently recur—highlighting promising avenues for future discovery.

#### Algorithm 1: Protein Structure Tokenization with Masks

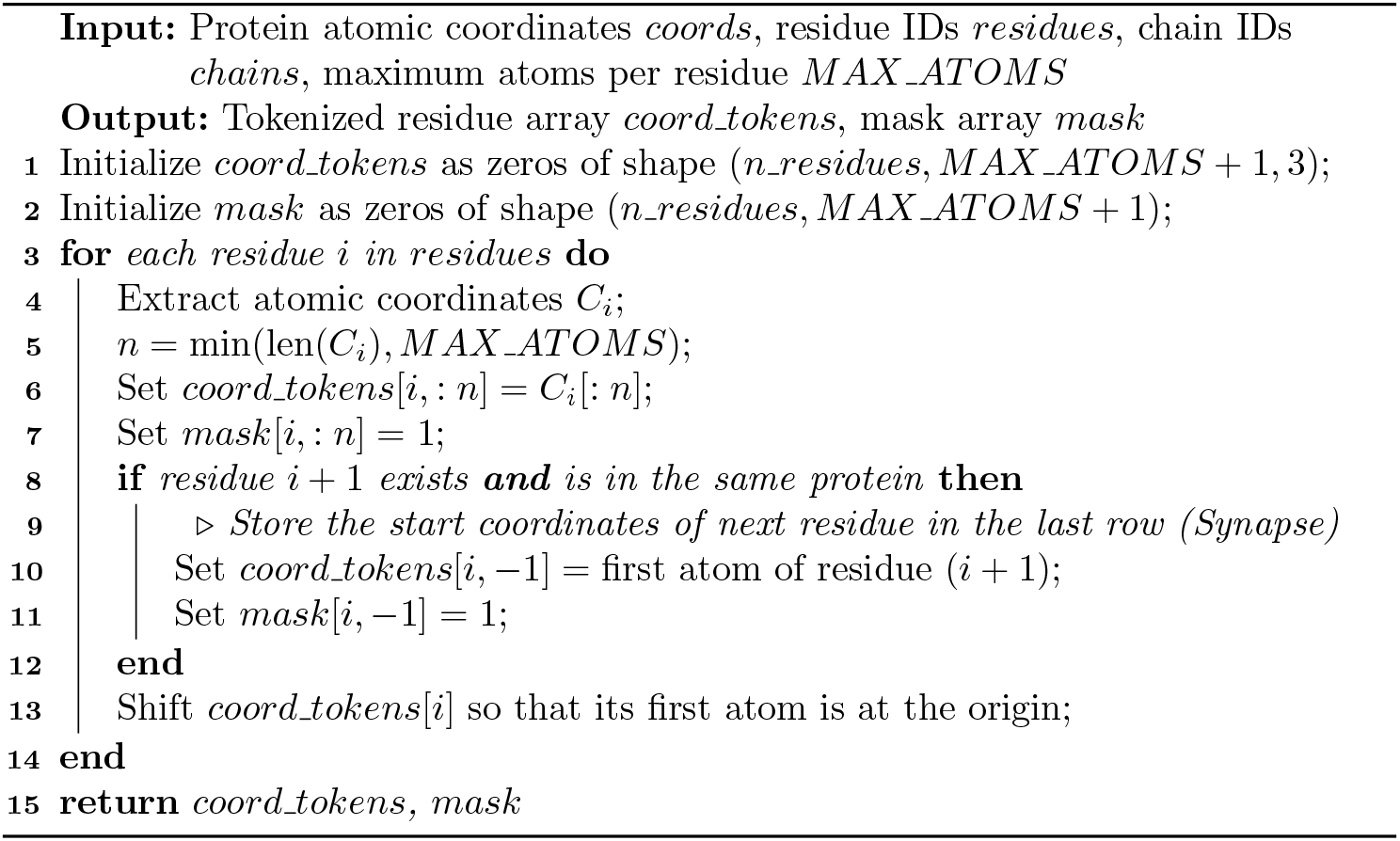

#### Algorithm 2: Protein Structure Detokenization with Masks

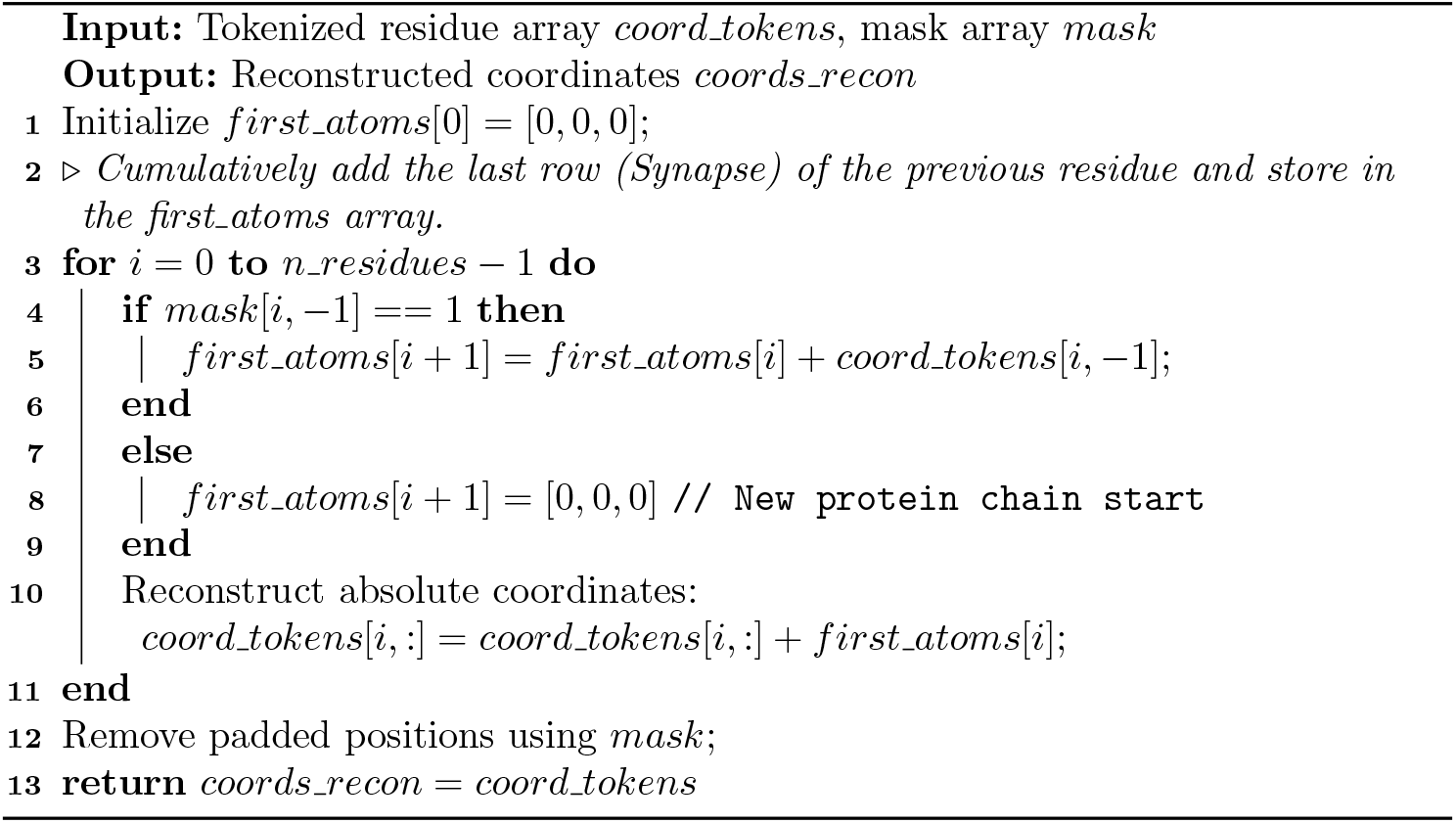

## 2. Results and Discussion

### 2.1 Method Overview

#### 2.1.1 Construction of Unfolded Protein Structure

ProteusFold translates an unfolded, idealized structure of each protein into its unique folded state, making the task inherently deterministic. Folded structures were obtained from the Protein Data Bank (PDB), with a focus on nuclear magnetic resonance (NMR) structures because their experimentally determined hydrogen positions enhance interpretability. The procedure for collecting these structures is detailed in Sec. S3.2.1, and the rationale for selecting NMR structures is provided in Sec. S1 of the Supplementary. The unfolded structures were constructed using PeptideBuilder [25], with uniform backbone dihedral angles corresponding to an extended chain conformation (Fig. 1.a) ^5^:

**Fig. 1:**
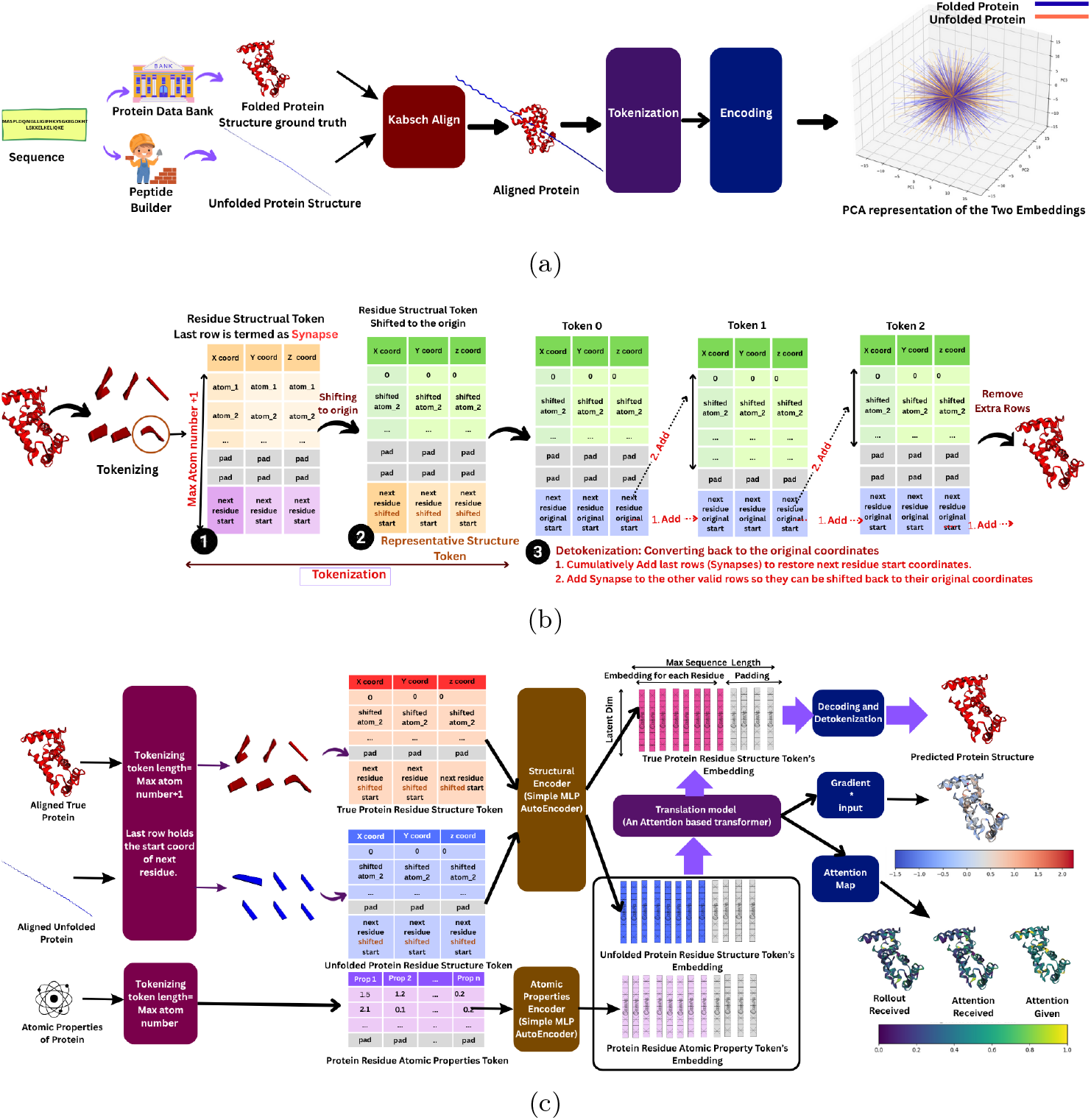
**(a)** Ground-truth protein structures are obtained from the PDB. For each protein, an unfolded structure is generated using `PeptideBuilder`. Both folded and unfolded structures are aligned using the Kabsch algorithm to remove unnecessary rotations and ensure standardization. The structures are then tokenized and encoded, with their PCA-reduced 3D representations visualized. The task is to translate between these two encodings. **(b)** Protein tokenization and detokenization method (illustrated here for the folded protein). **(c)** The complete workflow: tokenization, encoding, and translation to obtain the predicted protein structure. Additionally, protein characteristics are analyzed using attention and gradient × input attribution to understand residue-level contributions.

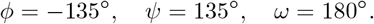

After construction, chain identifiers^6^ were reassigned to match the original PDB file, TER records were inserted and hydrogens were added using `PDBFixer` [26]. To standardize the start and target distributions, the Kabsch algorithm [27] is applied, and the folded coordinates are optimally superimposed onto the unfolded reference. This ensured both deterministic initialization and stable comparisons between folded and unfolded states.

#### 2.1.2 Structural Tokens and Atomic Properties

The core of this approach lies in *structural tokens*, which provides a standardized representation of the structure for each residue, with all its atoms, in both folded and unfolded states. As shown in Fig. 1.b^7^ and Alg. 1, each protein is split into residue-level tokens. Each structural token, denoted by coord tokens in Alg. 1, is a coordinate matrix 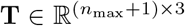, where *n*_max_ = 32 is the maximum number of atoms in any residue. For a residue with *n ≤ n*_max_ atoms, the first *n* rows of **T** contain its 3D atom coordinates, while the remaining *n*_max_ *− n* rows are filled with padding values (zeros) and accompanied by a binary mask that marks valid versus padded rows. The final row, called the *Synapse*, stores the starting coordinates of the next residue, ensuring bond connectivity across tokens. Finally, all valid coordinates (including the synapse) are shifted so that the first atom lies at the origin, yielding standardized structural tokens.

For *detokenization* (Fig. 1.b; Alg. 2), the Synapse rows are cumulatively summed to reconstruct the starting coordinates of subsequent residues. An array named first atoms is used to store these start coordinates, which are then applied to reposition each token and restore the global coordinates of all atoms. Finally, the Synapse and any padding rows are removed, yielding the full protein structure. This procedure preserves geometry and chain connectivity while enabling multi-chain proteins to capture protein–protein interactions.

For atomic properties, feature vectors 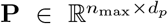 are extracted using the Mendeleev library [28], where *d*_*p*_ denotes the number of physicochemical descriptors. The extracted features include atomic number, weight, electronegativity, covalent and van der Waals radii, electron affinity, valency, density, polarizability, thermal conductivity, neutron count, specific heat capacity, and atomic radius. These atomic property tokens are then padded to *n*_max_ and normalized for subsequent processing.

#### 2.1.3 Embedding and Sequence Construction

Unfolded and folded structural tokens, as well as atomic property tokens, were encoded using compact multilayer perceptron (MLP) autoencoders (AE) [29] (Sec. S3.4), with a single AE for structural tokens and another for atomic properties. In both cases, the input is first flattened into 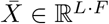, and the encoder maps it to a latent vector 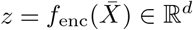 via *f*_enc_: ℝ^*L·F*^ *→* ℝ^*d*^, while the decoder reconstructs the input through *f*_dec_: ℝ^*d*^ *→* ℝ^*L·F*^. Specifically, structural tokens of size 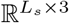 (*L*_*s*_ = *n*_*max*_ + 1 = 33) are flattened to 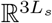 and encoded to ℝ^*d*^ (*d* = 128), and atomic property tokens of size 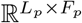 (*L*_*p*_ = *n*_*max*_ = 32, *F*_*p*_ = 13) are flattened to 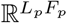 and mapped to ℝ^*d*^ (*d* = 128).

These structural and atomic property embeddings for each residue token are organized into fixed-length protein-level sequences for batch processing. Each sequence 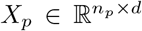 is truncated or zero-padded to a maximum length *L*_max_ = 400, with a binary mask 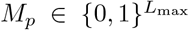 indicating valid residues. Stacking across *N* proteins produced tensors 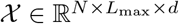 and 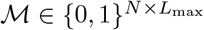. Using this approach, three parallel streams are constructed, namely atomic property embeddings (input), unfolded structural embeddings (input), and folded structural embeddings (target).

#### 2.1.4 Translation

As illustrated in Fig. 1.c, a transformer encoder–decoder architecture [30] is employed for sequence-to-sequence translation. Within the encoder, atomic property embeddings are processed using self-attention to capture inter-atomic relationships, producing a memory representation. The decoder operates on unfolded structure embeddings, where self-attention is applied to model intra-structural dependencies, and crossattention is performed over the encoder memory to integrate atomic-level information. The resulting folded structure embeddings are subsequently passed through the MLP decoder for structural tokens and finally detokenized to reconstruct full protein structures..

Note that this is an abridged description of the method; full methodological details are provided in Sec. S3 of the Supplementary.

### 2.2 Performance

#### 2.2.1 Brief discussion on Metrics

Protein structure prediction is commonly assessed using Root Mean Square Deviation (RMSD), which measures the average atomic deviation between predicted and reference structures [31]; Local Distance Difference Test (lDDT), which evaluates local residue-level accuracy [32]; Template Modeling score (TM-score), which quantifies global fold similarity [33]; Global Distance Test - Total Score (GDT-TS), which considers multiple distance thresholds to assess overall structural similarity [34]; and Predicted Aligned Error (PAE), which provides per-residue error estimates from AlphaFold predictions [6].

Protein-protein docking and interface quality are evaluated using DockQ, a composite score assessing the predicted interface [35]; Fraction of native contacts (Fnat), which measures how many native interface contacts are correctly recovered [36]; Ligand Root Mean Square Deviation (lRMSD), which measures deviation of the ligand chain in the complex [37]; and Interface Root Mean Square Deviation (iRMSD), which specifically evaluates structural deviation at the interface [37].

#### 2.2.2 Model’s Numeric Performance and Inference Time

The structural encoder–decoder MLP successfully reconstructed the original tokens for both folded and unfolded structures, achieving an *R*^2^ of 0.99997 and a mean squared error (MSE) of 0.0001201 on the test set, demonstrating its high fidelity in capturing structural features. Building on these embeddings, the translation model—trained for 570 epochs—achieved an MSE of 0.001052 and an *R*^2^ of 0.9994 on the test set, indicating excellent prediction of the target embeddings. In both cases, the models were selected based on their performance on the validation set.

Finally, a single 400-residue protein inference using ProteusFold takes approximately 0.0011 seconds, whereas ESMFold requires 14.2 seconds to predict the structure of a 384-residue protein on an NVIDIA V100 GPU [11]. In comparison, AlphaFold2 takes nearly six times longer than ESMFold for inference alone [11] and about ten minutes on average for end-to-end prediction [9].

#### 2.2.3 Protein Evaluation

Fig. 2.a–f summarizes per-chain and interface performance on the test set; panels (a)– (d) show violin plots for RMSD, lDDT, TM-score and GDT-TS stratified by chain length, panel (e) shows a histogram of DockQ for complexes, and panel (f) gives receptor-chain counts. Fig. 2a–d depicts the per-chain metrics for the test set protein sequences. The predictor attains very high per-chain accuracy for short and medium chains. For chains with length < 100 (*n* = 852), the mean RMSD is about 0.18 Å (median 0.17 Å), mean lDDT ≈ 0.998, mean TM-score ≈ 0.9995, and mean GDT-TS ≈ 99.995; the distributions are tight and concentrated near the ideal values (see the leftmost violins in each panel). For chains with 100 ≤ length < 300 (*n* = 393), the mean RMSD increases to ≈ 0.33 Å (median 0.30 Å) with slightly larger spread, mean lDDT ≈ 0.986, and mean TM-score ≈ 0.9992, indicating excellent global fold recovery but greater variability and a few outliers (RMSD up to ≈ 5.76 Å). Long chains (≥ 300, *n* = 4) show reduced local accuracy (mean RMSD 1.57 Å, mean lDDT ≈ 0.899, mean GDT-TS ≈ 78.2). The “all chains” violins (rightmost) summarize the aggregate behavior (*n* = 1249): mean RMSD 0.24 Å, mean lDDT ≈ 0.994, mean TM-score ≈ 0.9994, and mean GDT-TS ≈ 99.85. Overall, panels (a)–(d) indicate that the model reliably recovers native folds (high TM/GDT) and local geometry for short and medium chains, while long targets are the main source of elevated RMSD and reduced local scores.

**Fig. 2:**
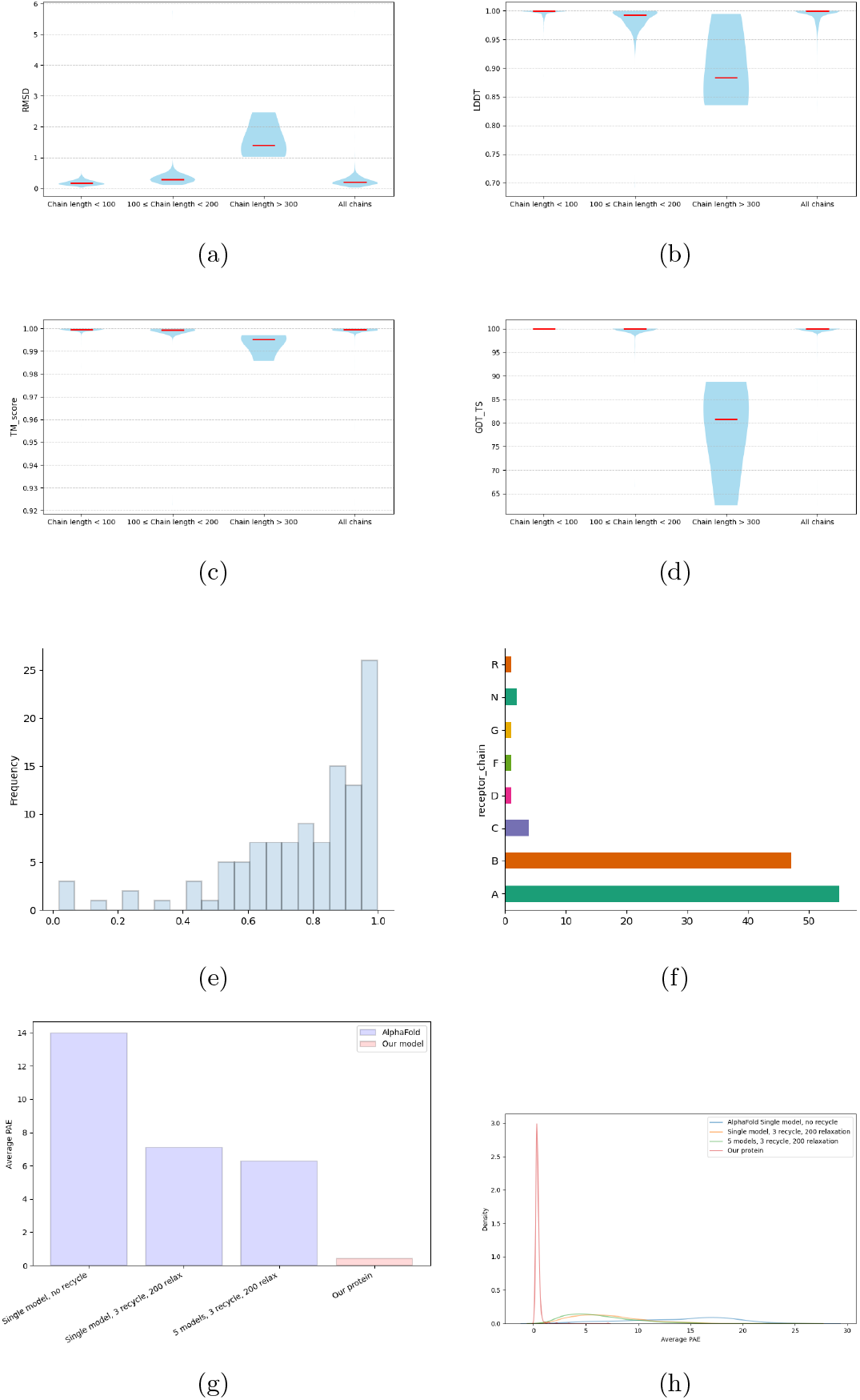
Performance summary across the test set. Panels (a)–(d) are violin plots (four violins: chain length < 100, 100 − 299, ≥ 300, all) for RMSD (Å), lDDT, TM-score and GDT-TS, respectively. Group means (medians) are: **(a)** RMSD — < 100: 0.18 (0.17) Å; 100–299: 0.33 (0.30) Å; ≥ 300: 1.57 (1.40) Å; all: 0.24 (0.20) Å. **(b)** lDDT — < 100: 0.998 (≈1.000); 100–299: 0.986 (0.992); ≥9 300: 0.899 (0.883); all: 0.994 (≈1.000). **(c)** TM-score — < 100: 0.9995 (0.9997); 100–299: 0.9992 (0.9995); ≥ 300: 0.9933 (0.9951); all: 0.9994 (0.9996). **(d)** GDT-TS — < 100: 99.995 (100.0); 100–299: 99.76 (100.0); ≥ 300: 78.22 (80.79); all: 99.85 (100.0). **(e)** Histogram of DockQ (complexes): mean = 0.7675, std = 0.2212, median = 0.8381; 95.5% of complexes have DockQ > 0.23; mean Fnat = 0.748, mean lRMSD = 2.795 Å, mean iRMSD = 1.747 Å. **(f)** Frequency of receptor assignments. **(g)** Mean avg-PAE (bars): AlphaFold2 (AF2) no-recycle = 13.97; AF2 3-recycle ≈ 7.10; AF2 5-model = 6.28; ProteusFold = 0.396 (AF2 in blue, ProteusFold (PF) in red). **(h)** KDE of avg-PAE: AF2 no-recycle (median 14.98, std 4.87), AF2 3-recycle (6.58, 3.34), AF2 5-model (5.63, 3.22), PF (0.357, 0.27). AF2 curves are broader and higher; PF is sharp near zero, showing lower and more consistent error.

Fig. 2.e shows the interface quality of the protein-protein complex in the test set. The DockQ histogram (n=112 complexes) shows strong interface performance: mean DockQ = 0.7675 (std = 0.2212), median = 0.8381, interquartile range approx. 0.656–0.939, with values spanning 0.018–0.998. Associated interface statistics are mean Fnat = 0.748, mean lRMSD = 2.795,Å and mean iRMSD = 1.747,Å. A large majority of complexes (107/112, 95.5%) exceed DockQ = 0.23, indicating that most predicted assemblies recover a meaningful fraction of native interfaces.

Fig. 2.f presents the frequency of receptor assignments. The horizontal bar graph illustrates how often each chain label served as the receptor in the test complexes, with chains labeled A and B appearing the most frequently. These counts reflect PDB labeling conventions rather than biological preference; hence, they are statistical insights and should not be interpreted as biologically meaningful. Finally, these panels collectively show that the model consistently produces near-native protein structures and accurately recovers interfaces across most complexes, as reflected by high DockQ and F_nat_ scores.

#### 2.2.4 Comparison with AlphaFold

As discussed in Sec. S3.1 of the Supplementary, limited computational resources prevented large-scale training of models such as AlphaFold or OpenFold from scratch. Instead, the pretrained ColabFold batch implementation of AlphaFold [12, 38] was employed to generate per-target predicted alignment error (PAE) values from the same FASTA inputs of the test set.^8^ Three AlphaFold configurations were evaluated in increasing order of complexity: (i) a single-model run with no recycling and no relaxation (fastest), (ii) a single-model run with three recycles and MMseqs2 with 200 relaxation steps, and (iii) the default AlphaFold ensemble (five-model ensemble, three recycles, MMseqs2 with 200 relaxation steps). The simple no-recycle configuration performed poorly, while recycling and ensembling improved PAE values. Nevertheless, across all configurations, AlphaFold performance remained substantially worse than that of ProteusFold on the evaluated targets.

Fig. 2.g presents vertical bars of the mean average PAE for the four methods (AlphaFold variants in blue, ProteusFold in red): AlphaFold (single-model, no recycle) = 13.97; AlphaFold (single-model, 3 recycles) = 7.10; AlphaFold (5-model ensemble, 3 recycles) = 6.28; and ProteusFold = 0.396. Fig. 2.h shows kernel density estimates (KDEs) of per-target average PAE for the same methods, highlighting the distributional spread (curve width reflecting variance). The AlphaFold curves are broader and centered at substantially higher PAE values (means ≈ 13.97, ≈ 7.10, ≈ 6.28; standard deviations ≈ 4.87, ≈ 3.34, ≈ 3.22 for the three respective configurations), whereas ProteusFold produces a sharply peaked distribution near zero (mean ≈ 0.396, standard deviation ≈ 0.27), reflecting both markedly lower and more consistent alignment error. Together, panels (g)–(h) demonstrate that ProteusFold achieves substantially tighter and lower PAE distributions compared with the tested AlphaFold variants, thereby indicating higher alignment confidence and local accuracy on this benchmark. It should also be noted that generating the AlphaFold results required approximately three days, whereas ProteusFold produced results within a few seconds (see Sec. 2.2.2 for a comparison of inference times). This stark contrast underscores the need for an open, lightweight model with strong performance capabilities.

### 2.3 Test Set Protein Examples

The visualizations of all proteins are compiled at [39]. For illustration, Fig. 3 shows selected examples of test set proteins alongside their predicted structures, highlighting the accuracy and structural fidelity of the model. Each row in the figure has three columns, where the first is the overlay of both predicted (blue) and reference (red) structures, the second is the predicted structure, and the third is the reference structure. In the following, a summary of the performance of the model for these proteins is presented, along with, where relevant, the quality of the predicted protein–protein interactions.

**Fig. 3:**
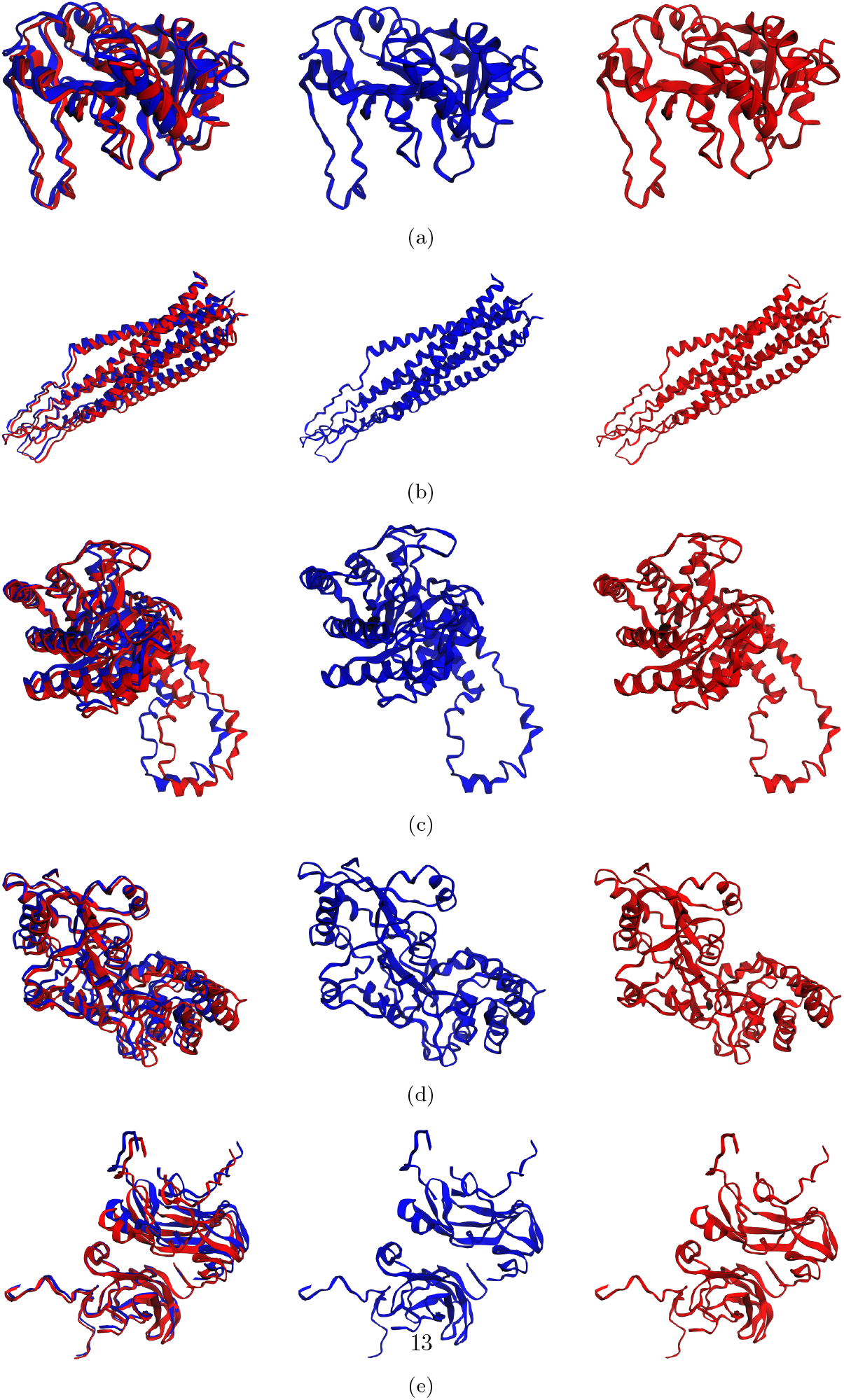
Comparison of predicted and experimentally determined protein structures. Each row corresponds to one protein; columns show (left) overlay, (center) predicted structure, and (right) true structure. **(a)** 1IEZ (Chain A): RMSD = 0.77 Å, GDT-TS = 94.55, TM-score = 0.998, lDDT = 0.906. **(b)** 2EZQ (Chains A–C): RMSD = 0.35–0.65 Å, GDT-TS = 97.79–100.00, TM-score ≈ 0.998–0.999, lDDT = 0.944–0.981. DockQ = 0.77, Fnat = 0.81, lRMSD = 2.23 Å, iRMSD = 1.28 Å. **(c)** 2GVE (Chain A): RMSD = 2.47 Å, GDT-TS = 62.59, TM-score = 0.986, lDDT = 0.835. **(d)** 1EZP (Chain A): RMSD = 1.68 Å, GDT-TS = 76.74, TM-score = 0.993, lDDT = 0.840. **(e)** 2G46 (Chains A–D): RMSD = 0.19–0.61 Å, GDT-TS = 98.96–100.00, TM-score ≈ 0.998–0.999, lDDT = 0.949–1.000. DockQ = 0.87, Fnat = 0.91, lRMSD = 2.32 Å, iRMSD = 0.79 Å.

#### 1IEZ (Chain A; length 217)

PDB ID 1IEZ represents 3,4-dihydroxy-2-butanone-4-phosphate synthase, a 47 kDa single-chain enzyme (217 residues) from *Escherichia coli* [40]. Fig. 3.a compares the predicted structure (second column) with the reference (third column), showing excellent agreement (RMSD = 0.77 Å, GDT-TS = 94.55, TM-score ≈ 0.998, lDDT = 0.906) and near-complete recovery of the native fold with very high global and local accuracy.

#### 2EZQ (Chains A–C; each 123 residues; total length = 369)

PDB ID 2EZQ corresponds to the trimeric ectodomain of simian immunodeficiency virus (SIV) gp41 (e-gp41), comprising three chains (A–C), each 123 residues long [41]. Fig. 3.b shows the prediction (second column), reference structure (third), and their overlay (first). Per-chain deviations are minimal (RMSD = 0.35–0.65 Å), with excellent agreement across GDT-TS (≈ 97.8–100), TM-score (≈ 0.998–0.999), and lDDT (0.944– 0.981). Interface evaluation further confirms accurate docking where Chain A serves as the receptor, while Chains B and C act as ligands. A DockQ of 0.77 and Fnat of 0.81 indicate a high fraction of native contacts recovered, while lRMSD = 2.23 Å and iRMSD = 1.28 Å demonstrate precise interface placement. Overall, both monomeric folds and assembly geometry are well captured for this trimeric complex.

#### 2GVE (Chain A; length 388)

PDB ID 2GVE represents a large single-chain (388 residues) D-xylose isomerase from *Streptomyces rubiginosus* [42]. Fig. 3.c shows the prediction (second column), reference (third), and their overlay (first). The predicted structure reproduces the overall fold (RMSD = 2.47 Å, TM-score = 0.986) but yields a lower GDT-TS (62.59) and moderate lDDT (0.835), indicating local deviations despite correct global topology. This indicates that although the global fold is accurately reproduced, local backbone placement for this protein exhibits some deviations.

#### 1EZP (Chain A; length 370)

PDB ID 1EZP corresponds to the 370-residue maltose-binding protein in complex with *β*-cyclodextrin [43]. Fig. 3.d shows a strong prediction with RMSD = 1.68 Å, GDT-TS = 76.74, TM-score = 0.993 and lDDT = 0.840. These metrics indicate accurate global fold recovery with reasonably good local geometry, reflecting reliable capture of the core and most secondary-structure elements for this large monomer.

#### 2G46 (Chains A–D; chain lengths = 119, 119, 20, 20; complex total length = 278)

2G46 is a four-chain assembly of the SET^9^ domain histone lysine methyltransferase (vSET) from *Paramecium bursaria* chlorella virus 1, bound to the cofactor S-adenosyl-*L*-homocysteine and a histone H3 peptide containing mono-methylated lysine 27 [44]. Fig. 3.e shows the prediction (second column), the experimental reference (third column), and their overlay (first column). Per-chain RMSDs are very small (0.19–0.61 Å), per-chain GDT-TS values are essentially saturated (98.96–100.00), and per-chain lDDT scores are high (0.949–1.000), confirming excellent local and global accuracy. Interface statistics, with Chain C as the receptor and Chains A, B, and D as ligands, are also strong: DockQ = 0.87, Fnat = 0.91, lRMSD = 2.32 Å, and iRMSD = 0.79 Å. These values indicate precise placement of interacting partners and faithful recovery of native interfacial contacts. Overall, the model successfully reconstructs both individual chain geometry and the higher-order assembly for this medium-sized complex.

Across these examples, the model reliably recovers native folds (high TM-scores) and shows particularly strong accuracy for smaller chains and tightly packed complexes (low per-chain RMSD, high lDDT). Multi-chain targets (2EZQ, 2G46) additionally exhibit robust interface recovery as measured by DockQ and Fnat, whereas very long single chains (e.g., 2GVE, length 388) may still display elevated local deviations despite correct global topology.

### 2.4 Residue-Level Analysis

Two types of interpretation information were collected for residue-level analysis, namely the attention [30] maps from the Transformer and the gradient-based feature attribution using *gradient × input* [45, 46]. The method of collecting attention has been shown in Sec. S4.1 in the Supplementary. Briefly, the attention matrix from the encoder is used for interpretation as they contain the atomic information of each residue, and that is the most biologically meaningful. Using the matrix, three values for each residue were obtained namely, “attention-recieved” (column-wise summation of the matrix), “rollout-recieved” (column-wise normalized sum), and finally “attention-give” (row-wise summation). All three give different perspectives on the same information. It is adivised to mainly focus on the matrix and use the three derived values for nuances. These attention maps can be interpreted as indicators of global structural importance, where residues that consistently receive high attention are more informative and influential to the overall protein structure. These maps reveal dependency networks among residues, highlighting key anchoring contacts, cooperative stabilization pathways, and potential targets for experimental mutational or kinetic investigations.

Gradient-based attribution reflects how input features influence a selected target, indicating whether they increase (positive attribution) or decrease (negative attribution) its value. For a target such as the predicted structure *y*_pred_, this is expressed as *∇*_*x*_*y*_pred_ *⊙ x*, where *x* denotes the input features. However, when the target itself is a protein structure, the notions of “increase” or “decrease” are not meaningful, making direct attribution difficult to interpret. To address this, the mean squared error (MSE) loss was instead chosen as the attribution target, yielding *∇*_*x*_ ((*y −y*_pred_)^2^)*⊙ x*, which can be interpreted as measuring how each residue in the input sequence influences the model’s tendency to move closer to the native structure (by decreasing MSE) or farther away (by increasing MSE). A negative attribution value at residue *i* indicates that the wild-type identity at *i* contributes to producing a more native-like coordinate for that position (i.e., it reduces predicted per-residue error / RMSD relative to the model baseline). Conversely, a positive attribution at *i* indicates that the residue’s identity drives the model away from the native coordinate (increases predicted error). Thus negative attribution can be interpreted as model-inferred stabilizing/local-native contribution; positive attribution can be taken as model-inferred destabilizing/local-nonnative contribution. A detailed description and interpretation of the residue-level gradient-based attribution is provided in Sec. S4.2, and the generalized interpretation using *gradient × input* with the mean squared error (MSE) as the loss function is presented in Sec. S6.

Fig. 4 illustrates these two types of diagnostics for two example proteins: Protein G B1 domain (PDB: 1GB1) and Protein Nucleocapsid Zinc Fingers (PDB: 1AAF). The first four rows of the mentioned figure presents the visualizations for Attention machanism analysis for both the protein (1GB1 left and 1AAF right):

**Fig. 4:**
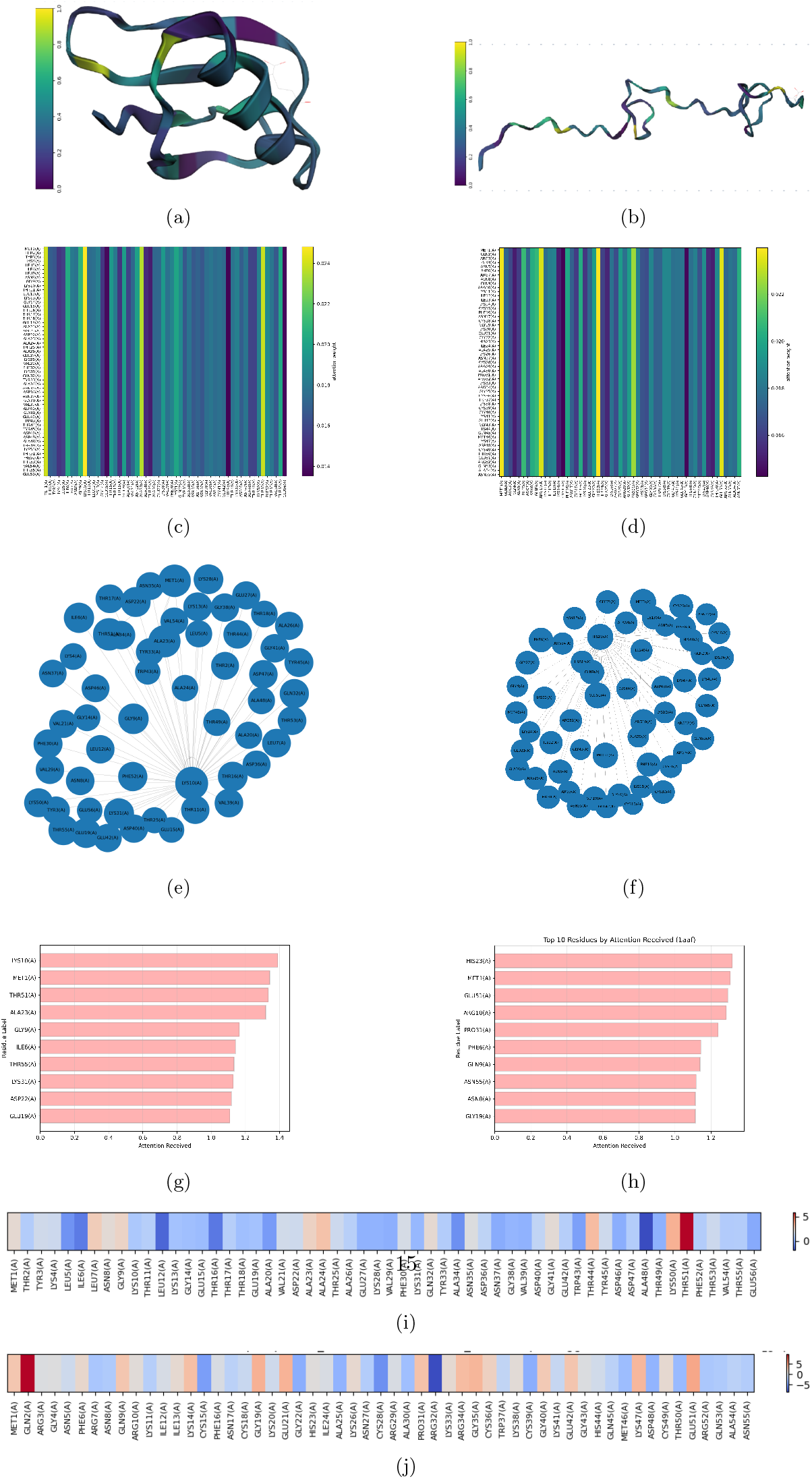
Rows 1–4 show Attention diagnostics for Protein G B1 (PDB: 1GB1, left column) and 1AAF (right column). Row 1-4: (a,b) per-residue attention on 3D structures; (c,d) attention matrices; (e,f) residue–residue attention graphs; (g,h) top-10 residues by received normalized attention. (i) *gradient×input* heat map for 1GB1; (j) *gradient×input* heat map for 1AAF. Attention highlights structural anchors; *gradient×input* reflects kinetic influence.

- Row 1 (Fig. 4.a,b) — 3D renderings of per-residue “attention-recieved”, obtained by column-wise summation of the attention matrix (see Sec. S4.1). The values are color-mapped onto the protein structure (see Supplementary Fig. S2 for additional visualizations such as “rollout-recieved” and “attention-given” maps). The actual figures are interactive 3D structures that can be used as tools to explore and interpret residue-level attention in greater detail.
- Row 2 (Fig. 4c,d) — the attention matrices for each protein (pairwise attention between residues).
- Row 3 (Fig. 4.e,f) — attention graphs that summarize residue–residue dependency networks inferred from the matrices. This map helps to understand the relationships between each residues and their influences as their size is dependant on their importance.
- Row 4 (Fig. 4.g,h) — the top ten residues ranked by total “rollout-received” i.e. normalized attention.

Finally, Fig. 4.j and Fig. 4.k show the *gradient× input* heat maps for 1GB1 and 1AAF, respectively, allowing direct comparison of individual residue influence. The analyses for these two proteins are discussed below. It is also advised to refer to Table S3 for the three-letter and one-letter representations of each residue, as the following analysis alternates between full names, three-letter codes, and one-letter codes.

#### 2.4.1 Protein G B1 domain (1GB1)

Protein G B1 domain (1GB1) is a small, well-characterized globular protein of 56 amino acids, derived from the immunoglobulin-binding protein G of *Streptococcus* [47]. Its simple structure—a four-stranded *β*-sheet packed against a single *α*-helix—makes it a widely used model system for studying protein folding, stability, and design [48– 52]. Using Fig. 4.a, c, e, and g, the attention mechanism for this protein was analyzed, while Fig. 4.i shows the heatmap from the *gradient × input* analysis. The top residues identified by both analyses are presented below.

##### Attention Analysis of Lysine-10 (LYS10/K10)

In ProteusFold, Lysine-10 receives the highest attention score among 1GB1 residues, suggesting that it functions as a structural anchor whose state strongly influences multiple other residues, a finding that is consistent with prior experimental data [48–51]. Specifically, Olson *et al*. [48] showed that mutations in the dynamic loop (residues 9–12) produce disproportionate epistasis effects, thereby underscoring its functional importance. In line with this, structural analysis revealed that this loop is coupled to the *β*_1_–*β*_2_ loop via hydrogen bonds between Glu56 and the Asp40/Lys10 amides, with additional IgG-Fc^10^ contacts at Val39. Collectively, these observations highlight Lys10 as a key linkage point between structural elements essential for folding, stability, and function.

Moreover, McCallister *et al*. [49] demonstrated that mutations in the first *β*-turn (e.g., a six-Gly insertion after Lys-10, “K10+6G”, or T11A) destabilize GB1 while leaving the folding rate largely unchanged. For K10+6G, they reported ΔΔ*G ≈* 1.42 kcal, mol^−1^ with a low Φ-value (Φ ≈ 0.21), which is consistent with the turn forming after the rate-limiting transition state ^11^ and contributing mainly to equilibrium stability rather than *k*_*f*_. Further supporting this, molecular dynamics studies [50] suggest that breakage of the E56–K10 salt bridge can trigger *β*-hairpin disruption, indicating that K10 functions as an anchoring contact whose loss promotes unfolding. Finally, structural analyses likewise confirm a persistent E56–K10 interaction in the native ensemble [51].

Taken together, the model’s attention signal and independent experimental and simulation evidence strengthen the interpretation of K10 as a *structural anchor*, nominating it as a high-priority target for future studies.

##### Attention and Gradient × Input Analysis of Threonine-51 (Thr51/T51)

Thr51 was found to be the third-highest attention-receiving residue and to exhibit the most positive (MSE-increasing) *gradient × input* value in the model, indicating that it conveys substantial information about other positions while contributing a destabilizing or locally nonnative effect. Consistent with this, Kuszewski *et al*. [53] report a site-resolved unfolding/exchange rate

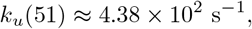

indicating rapid local exchange. This kinetic lability causes T51 to frequently leave its native hydrogen-bonded register, allowing it to report on folding–unfolding transitions; as a result, attention mechanisms prioritize it because its instantaneous state provides information that reduces uncertainty about correlated structural changes. Overall, the concordance between the high experimental *k*_*u*_ and the model’s elevated attention score, together with a strongly positive *gradient × input* signal (indicating increased MSE), suggests that T51 is highly informative for other residues and contributes to structural destabilization, warranting further experimental validation.

##### Gradient × Input Analysis of Alanine-48 (Ala48/A48)

The model exhibits a high negative *gradient × input* for Ala48, which aligns with prior research. Specifically, studies indicate that Ala48 is highly solvent-exposed and forms part of an amphiphilic helix, where its side chain packing contributes to the stabilization of local structure and folding intermediates [47]. Furthermore, experimental mutation studies by McCallister *et al*. [49] demonstrated that replacing Asp46 disrupts its hydrogen bond with Ala48, thereby causing a 20-fold decrease in folding rate without affecting unfolding. Taken together, these observations highlight Ala48’s crucial role in stabilizing the folding transition state through this interaction.

#### 2.4.2 Protein Nucleocapsid Zinc Fingers (1AAF)

The HIV-1 nucleocapsid protein (PDB: 1AAF) contains two highly conserved CCHC-type zinc finger motifs that play essential roles in viral RNA recognition, genome packaging, and reverse transcription [54, 55]. These compact zinc-binding domains provide an experimentally tractable model for studying zinc finger structure, folding, and nucleic acid interactions. Using Fig. 4.b, d, f, and h, the attention mechanism for this protein was analyzed, while Fig. 4.j shows the heatmap from the *gradient × input* analysis. The top residues identified by both analyses are presented below.

##### Attention Analysis of Histidine-23 (His23/H23)

As shown in Fig. 4.d and Fig. 4.g, His23 receives the maximum attention score in the protein 1AAF. It is interpreted as strong evidence for both structural and functional importance of the referred residue, supported by independent biochemical and mutational studies, as shown in the following.

His23, the histidine of the first CCHC zinc-finger motif (Cys–Cys–His–Cys), directly coordinates Zn^2+^, thereby stabilizing the local fold of the first finger domain [52, 56].

Furthermore, as a key residue at the structural core of the nucleocapsid’s retroviral-type zinc finger [56], His23 is indispensable for both the domain’s structural integrity and its functional activity. Consistent with this, site-directed mutagenesis, such as H23C^12^, disrupted the ^1^H NMR-derived structure and, noteably, abolished viral infectivity [57, 58]. Mechanistically, replacing the imidazole ligand with a thiol perturbs zinc coordination geometry, destabilizes the zinc knuckle, impairs nucleic-acid chaperone activity, and consequently compromises HIV-1 replication [52, 59]. Taken together, the strong model attention on His23, its established role as a Zn^2+^ ligand, and the clear loss-of-function phenotypes observed upon mutation support the conclusion that this residue is critical for structural stability and HIV-1 infectivity. Therefore, His23 represents a high-priority candidate for follow-up studies using spectroscopy, mutagenesis, or inhibitor design.

##### Gradient × Input analysis of Glutamine-2 (Gln2/Q2) and Arginine-32 (Arg32/R32)

Gradient-based analysis revealed that Gln2 shows a strong positive gradient attribution, indicating an increasing role in MSE, while Arg32 exhibits a strong negative attribution, indicating a decreasing role in MSE. As discussed earlier, this suggests that Arg32 acts as a stabilizing residue, whereas Gln2 contributes to destabilization. The following discussion provides further support for this interpretation.

Mimic study by Druillennec *et al*. show that this complex is stabilized by hydrophobic contacts of residues belonging to both zinc fingers and ionic interactions of Arg-26 and Arg-32 with the ribose phosphate backbone [60]. The stabilizing role of this region has also been supported by Morellet *et al*. [52] and Summers *et al*. [55].

For the case of Gln2, since no direct mutational studies on residue Gln2 (Q2) could be found, DynaMut [61] was used to explore its potential role in the folding process. Although stability predictors provided mixed ΔΔG estimates for Q2 substitutions, Elastic Network Contact Model (ENCoM)[62] consistently indicated increased vibrational entropy, suggesting enhanced local flexibility at this N-terminal position. This indicates that Q2 may act as a labile site, affecting local dynamics and contributing to localized destabilization. Verification of this hypothesis would require site-resolved Hydrogen–deuterium (H–D) exchange [63] or NMR relaxation [64] experiments to assess whether Q2 indeed exhibits elevated unfolding kinetics (*k*_*u*_).

### 2.5 Atomic-Level Analysis

At the atomic-level, *Gradient × Input* values for the properties defined in Sec. S3.3.2 (Supplementary) were computed using the procedures described in Sec. S4.3 (Supplementary). Briefly, for each residue *r*, the residue embedding **e**_*r*_ received a gradient signal 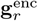 from the supervised loss (Eq. S10). These gradients were then backpropagated through the atomic encoder to assign contributions to each atomic property. A simplified attribution for atomic property *x*_*r,a,p*_ can be expressed as

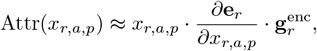

where *r, a*, and *p* index residues, atoms, and atomic properties. Here, 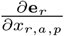 is the gradient of the residue embedding with respect to the atomic property, and the element-wise product with 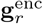 propagates the residue-level importance to the atomic-level. This is a simplified and intuitive version of the full vector–Jacobian product (VJP) [65, 66] in Eq. S13.

Fig. 5.a shows the collective (absolute sum) *gradient × input* values for each property, grouped by element and ordered by the total contribution of each property.

**Fig. 5:**
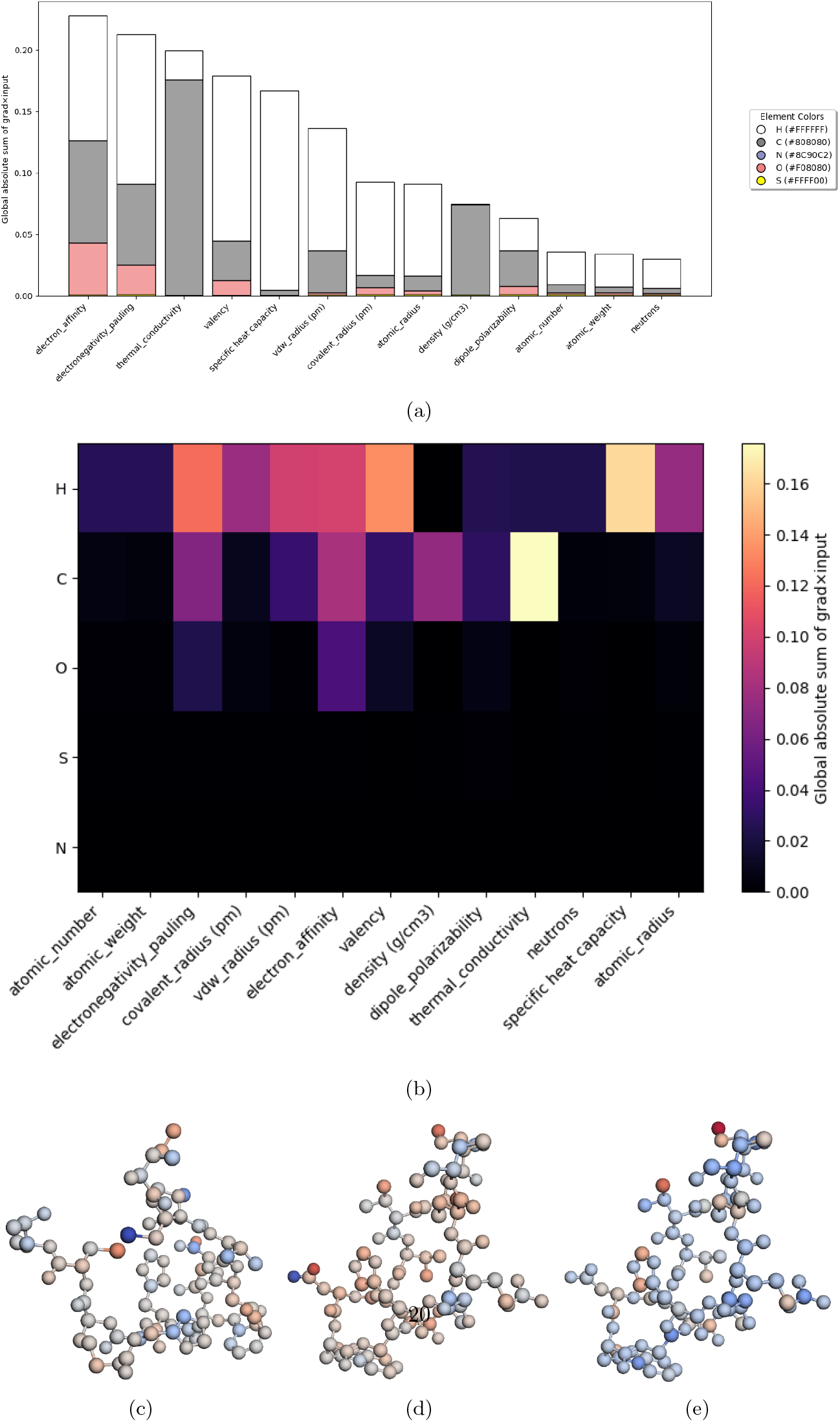
**a**. Histogram of the element-wise absolute sum of *gradient × input* values for atomic properties in protein folding. **b**. Heatmap showing the absolute summed contributions of each element across all atomic properties. **c–e**. Interactive structures of Compstatin (1A1P) colored by *gradient × input* values for (**c**) electronegativity, (**d**) electron affinity, and (**e**) thermal conductivity. Blue indicates more negative values (decreasing MSE), while red indicates more positive values (increasing MSE).

From this figure, it becomes clear that the interplay of hydrogen, carbon, and oxygen electronegativity has a pronounced impact on the MSE of protein folding. This observation is consistent with prior studies showing that differences in electronegativity generate partial charges and polar bonds, which in turn drive hydrogen bonds, dipole interactions, and salt bridges that shape folding energetics and stability [67, 68]. In addition, electronegative clusters or acidic regions are known to further modulate folding propensity [69]. Furthermore, hydrophobicity—another fundamental driving force in protein folding [70]—arises from low bond polarity. Specifically, nonpolar C–H bonds promote the burial of hydrocarbons, whereas polar O–H and N–H bonds remain solvent-exposed and participate in stabilizing interactions [70–76]. Values for these properties across the relevant elements can be cross-checked in Table S1 of the Supplementary.

It can also be observed that oxygen exerts a particularly strong effect through its electron affinity and electronegativity. Oxygen’s high electron affinity and electronegativity critically determine hydrogen bond acceptor strength, dipole interactions, and solvent effects, which are key to maintaining the directional hydrogen-bonding network essential for folding specificity and stability [67, 77, 78]. Moreover, oxygen’s moderately high values observed in these properties help explain why electronegativity and electron affinity collectively ranked higher than the thermal properties (thermal conductivity and specific heat), even though the contributions of individual elements were not the largest.

From Fig. 5.a and Fig. 5.b, it is further evident that the thermal conductivity of carbon has the highest individual element importance, underscoring its crucial role in the folding process. This interpretation is reinforced by atomic-resolution experiments highlighting the significance of carbon’s thermal conductivity in folding [79]. Similarly, computational studies have quantified the thermal conductivity of proteins at the atomic level, reporting values consistent with experimental data and showing that heat transfer along the main chain is dominated by local vibrations, particularly those involving carbon atoms in polypeptide bonds [80]. Furthermore, Fig. 5 indicates that the specific heat of hydrogen strongly influences the MSE of folding prediction. This observation is consistent with protein folding thermodynamics, where heat capacity changes (Δ*C*_*p*_) arise from both hydrophobic effects and hydrogen-bond networks. In this context, Cooper [81] proposed that Δ*C*_*p*_ reflects the cooperative ordering and melting of hydrogen bonds in folded proteins, with hydrogen atoms central to these interactions. Experimental studies further support this interpretation: Mallamace *et al*. [82] linked temperature-dependent hydrogen bond lifetimes to Δ*C*_*p*_ variations, while Loladze *et al*. [83] emphasized the role of buried polar groups in stabilizing proteins through heat capacity changes. Taken together, this discussion demonstrates that hydrogen and carbon collectively exert a significant influence on folding. However, as the following analysis will show, not all atoms exert equally strong effects, and the contributions of individual atom elements often differ markedly from their collective impact.

Fig. S3.a (Supplementary) presents a pie chart showing the proportion of each element in proteins. Hydrogen and carbon dominate, while oxygen and nitrogen make up a modest portion, and sulfur is present in very small amounts. However, as shown in Fig. S3.b and Fig. S4 of the Supplementary, not all hydrogen or carbon atoms have a large impact on folding despite their abundance. In contrast, sulfur, although much less common, can have a substantial effect on protein folding. This is illustrated in Fig. S3.b, which shows that the small proportion of sulfur atoms exerts a disproportionate influence across multiple properties. Fig. S3.b and Fig. S4.a show sulfur’s dominance in most properties, marked by high variability in electron affinity and polarizability, and notable effects on radii and electronegativity. Its large size and polarizability facilitate unique stabilizing interactions in proteins (see Tables S1, S2), with methionine and cysteine contributing via disulfide bonds and chalcogen bonding [84–87]. This is further reinforced by divalent sulfur’s ability to form both chalcogen bonds through its *σ*-holes and hydrogen bonds via lone pairs [86, 87], stabilizing secondary structures through helix capping, *β*-sheet edge protection, and *β*-turn augmentation. The strength of such chalcogen bonds can rival hydrogen bonds, underscoring sulfur’s outsized role in folding beyond its relative abundance. Further discussion are conducted in Sec. S7 in the Supplementary.

Now that the effects of each element on the various atomic properties have been established, and it has also been shown that not all atoms of a given element exert the same importance, it becomes necessary to identify which specific atoms are most relevant for each atomic property in the protein under investigation. A demonstration of the atomic-level analysis process is provided below using Compstatin (PDB: 1A1P) as an example case.

#### 2.5.1 Atomic Analysis of Compstatin (1A1P)

Compstatin (1A1P) is a protein that binds specifically to Complement component 3 (C3) ^13^ and inhibits complement activation [89]. Fig. 5.c shows the atomic visualization of the gradient × input of electronegativity. Upon analysis, it was found that the most negative (blue) atom, that is, the one that helps reduce the MSE for this property, is the sulfur at position 5 in Cys2. Following this, the next most negative (blue) atoms are primarily hydrogens and carbons from residues such as Val3, His10, and others. Fig. 5.d presents the gradient × input values for electron affinity, showing that the oxygen at position 7 in Asp6 has the largest negative impact on MSE, followed by hydrogens from various residues. For thermal conductivity, Fig. 5.e shows that the negative impacts are primarily dominated by carbon, with the carbon at position 7 of His10 having the strongest effect. Although atomic-level experiments specifically targeting Compstatin (1A1P) are lacking, existing molecular dynamics simulations and experimental binding studies indicate that electronegativity plays a significant direct role [90], electron affinity influences folding indirectly through hydrogen-bond donor/acceptor properties [91, 92], and thermal conductivity contributes via thermal motion effects [90]. This suggests ample room for further atomic-level exploration in this area.

Overall, the mentioned findings provide significant insight into the driving factors of protein folding, highlighting the interplay between structural encoding and intrinsic atomic properties. It also highlights the practical use of ProteusFold for conducting analysis on each protein in an atomic-level.

### 2.6 Positional analysis and hypothesis of folding hotspots

A large body of research indicates that protein folding is not equally sensitive across all residues, but is instead dominated by hotspot regions that recur across diverse folds. Meta-analyses of *ϕ*-value data and comparative studies across many unrelated proteins show that residues in N- or C-terminal regions, short loops (particularly *β*-hairpins), and helix-capping motifs are disproportionately represented in experimentally identified folding nuclei [93–95]. Residues within the hydrophobic core are broadly critical, as mutations at buried positions frequently perturb folding transition states and alter folding kinetics across diverse protein folds [96, 97]. Taken together, these findings indicate that certain topological locations -termini, loops/turns, and hydrophobic nuclei - have a substantially higher probability of containing influencing residues for protein folding than other regions.

Fig. 6 presents the positional analyses of residue importance scores. Fig. 6.a shows the residue positions that receive the highest attention and rollout values; clear peaks are visible, notably near residue IDs 1, 10, 23, 27, and 51, with smaller spikes at around residue IDs 100 and 200. For further analysis, the procedure described in Sec. S5 of the Supplementary was applied. Fig. 6.b shows the locations of interest identified by that method, which cluster around the peaks observed in panel (a).

**Fig. 6:**
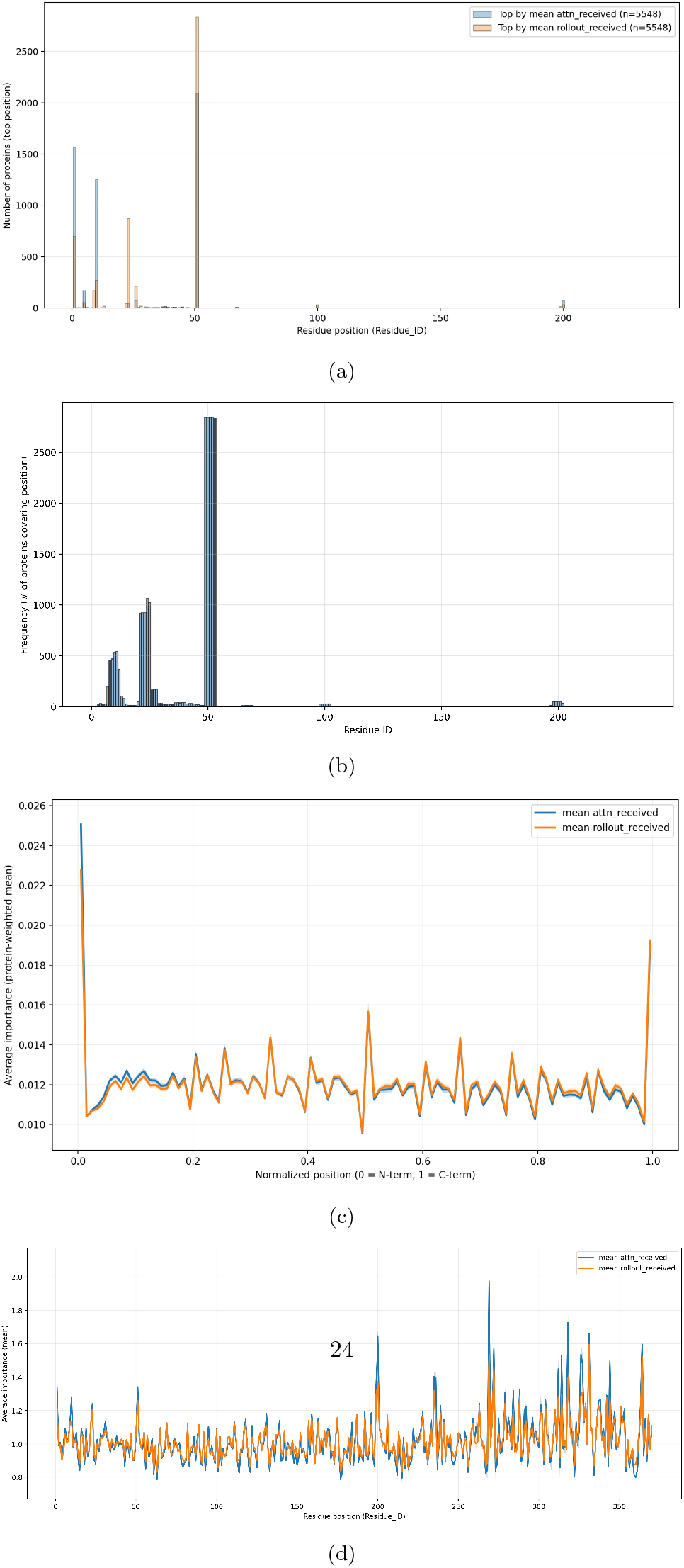
Positional analysis of residue importance. (a) highest attention and rollout recieving residues. (b) Locations of interest (identified by the method in Sec. S5 in Supplementary) coverage across the protein. (c) Normalized position analysis of the average attention showing enrichment near the sequence midpoint (0.5). (d) Average attention on the absolute residue IDs. It is evident from these figures that specific regions consistently harbor important residues, referred to as *folding hotspots*.

To account for proteins of varying lengths, the average per-protein normalized attention was computed across normalized residue positions and is shown in Fig. 6.c. In this analysis, elevated importance is revealed in the central region of proteins, with a pronounced increase near the normalized position 0.5. The average normalized importance mapped back to actual residue IDs is shown in Fig. 6.d. The results are largely consistent with the previous discussion, although several pronounced peaks at residue IDs beyond 250 are additionally observed. These extreme peaks are attributed to sparse sampling of very long proteins, which can inflate the mean; increasing the dataset size is expected to smooth these artifacts.

Overall, it is indicated by these results that certain sequence regions are more likely to contain high-importance residues. Based on these observations, it is hypothesized that functionally and structurally important residues are preferentially placed by nature in particular regions of proteins; such regions are referred to as *folding hotspots*. This constitutes a testable hypothesis, and further targeted experiments and larger-scale analyses will be required to confirm or refute it.

### 2.7 Future Research

This study focuses exclusively on pure proteins and protein–protein complexes, as PeptideBuilder currently supports only pure protein structures. Future research should explore the development of tools capable of modeling proteins in combination with other biomolecules, such as DNA and RNA, thereby broadening the scope of structural studies. Researchers possessing substantial computational resources could construct more powerful models with larger context windows, enabling the prediction and analysis of very long proteins and their folding mechanisms. Another promising direction is the systematic collection of misfolded proteins associated with various diseases and their analysis at both the residue and atomic-levels to uncover the underlying causes of misfolding using ProteusFold. In addition, large-scale studies examining the importance of proteins across specific regions may help validate the proposed “*folding hotspot* “ hypothesis. Finally, advances in encoding strategies could substantially reduce memory requirements, improving efficiency and accelerating progress in protein structure prediction.

## 3 Conclusion

ProteusFold demonstrates strong capabilities in predicting protein structures directly from sequence. Protein–protein interactions within complexes are also effectively captured, while the model remains lightweight enough for practical applications. A key contribution is the novel protein structure tokenization scheme, which not only simplifies the problem but also preserves residue connectivity across chains within the same protein. This framework further enabled residue-level explanations, many of which were consistent with prior experimental findings. Such consistency increases confidence that the interpretability approach can serve as a valuable starting point for experimental studies on novel or less-characterized proteins. Moreover, ProteusFold provides, to the best of current knowledge, the first atomic-level insights into properties and atoms critical for protein folding. Additionally, positional analysis suggests the potential existence of protein “*folding hotspots*,” which may represent a promising avenue for future research. Taken together, these advances establish ProteusFold as a significant step forward in protein folding research, bringing the field closer to a deeper understanding of the underlying biology.

## Declarations

### Data availability

All data can be found at the following link:https://www.kaggle.com/datasets/sakibahmed91/protein-data

The original PDB files of the folded structures have been provided, together with the converted CSV files of the folded and unfolded structures. The combined CSV file containing both folded and unfolded structures used for training and testing has also been included.

### Code availability

All code can be found at the following link: https://github.com/bojack-horseman91/ProteusFold

### Author contribution

**Saleh Sakib Ahmed:** Conceptualization, Data Collection, Method Development, Analysis, Manuscript Writing, Manuscript Editing and Reviewing

## Supplementary

**Table S1:** Electronegativity, electron affinity, and thermal conductivity of selected elements.

| Element | Electronegativity(Pauling) | Electron Affinity(kJ/mol) | Thermal Conductivity(W/m·K) |
| --- | --- | --- | --- |
| Hydrogen (H) | 2.20 | 0.754 | 0.1815 |
| Carbon (C) | 2.55 | 1.262 | 1.59 |
| Nitrogen (N) | 3.04 | -1.40 | 0.026 |
| Oxygen (O) | 3.44 | 1.461 | 0.027 |
| Sulfur (S) | 2.58 | 2.077 | 0.27 |

**Table S2:**
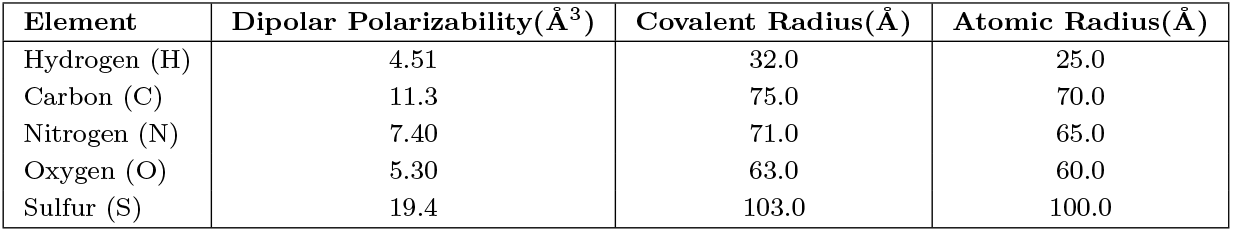
Dipolar polarizability, covalent radius, and atomic radius of selected elements.

| Element | Dipolar Polarizability($\text{\AA}^3$) | Covalent Radius($\text{\AA}$) | Atomic Radius($\text{\AA}$) |
| --- | --- | --- | --- |
| Hydrogen (H) | 4.51 | 32.0 | 25.0 |
| Carbon (C) | 11.3 | 75.0 | 70.0 |
| Nitrogen (N) | 7.40 | 71.0 | 65.0 |
| Oxygen (O) | 5.30 | 63.0 | 60.0 |
| Sulfur (S) | 19.4 | 103.0 | 100.0 |

### S1 Discussion on Protein Structure Experiments and Importance of Hydrogen Bonds

Hydrogen atoms are crucial for understanding protein structure and dynamics, yet their experimental detection is non-trivial. Most structural biology techniques rely on scattering by electrons or nuclei, and since hydrogen has only one electron, it is often invisible to standard approaches.

#### S1.1 Experimental Detection of Hydrogens

Neutron diffraction can directly visualize hydrogens (or deuterium-substituted atoms) because neutrons interact with atomic nuclei rather than electron clouds [98–100]. However, neutron crystallography requires very large, well-ordered crystals and access to neutron sources, limiting its widespread use. X-ray crystallography, the dominant method for high-resolution protein structures, typically does not resolve hydrogen atoms because of their weak scattering. Only at ultra-high resolutions (≲ 1.0 Å) do some hydrogens become visible in electron density maps. For direct and unambiguous localization of hydrogen (or deuterium) atoms, neutron crystallography or joint neutron–X-ray refinement is used [101, 102]. Cryo-electron microscopy has recently achieved near-atomic resolutions, but even at 1.2 Å maps, hydrogens are generally not reliably observed [103–105]. In contrast, Nuclear Magnetic Resonance (NMR) spectroscopy directly measures signals from hydrogen nuclei. Through ^1^H chemical shifts, nuclear Overhauser effects (NOEs), and scalar couplings, NMR provides detailed, solution-state information on hydrogen positions and interactions [106– 109]. Thus, among routine experimental techniques, only NMR consistently gives hydrogen-specific information.

#### S1.2 Importance of Hydrogen Bonds in Protein Folding

Hydrogen bonds are fundamental stabilizing forces in protein folding. They guide the formation of secondary structures, such as *α*-helices and *β*-sheets, and help establish precise tertiary arrangements by stabilizing side-chain and backbone interactions [31]. The cooperative network of hydrogen bonds reduces conformational entropy and drives the protein into its unique native state. Disruption of hydrogen bonding patterns often leads to misfolding, aggregation, or loss of function [110, 111].

Because NMR directly provides information on hydrogen atoms, it is uniquely suited to characterize hydrogen-bonding networks in proteins under physiological conditions. Moreover, since hydrogen bonds play a key role in protein folding, NMR structures are therefore ideal as reference structures for building interpretable models that can accurately capture the folding mechanism.

### S2 Evaluation Metrics

To evaluate the performance of protein structure prediction, several widely used metrics are employed. The Root Mean Square Deviation (RMSD) measures the average atomic distance between a predicted and reference structure after optimal superposition, typically ranging from 0 Å upward, where values below 2 Å indicate near-native accuracy [31]. The Local Distance Difference Test (lDDT) provides a superposition-free score by comparing interatomic distances, with scores between 0 and 1 (higher values indicating more accurate models) [32]. The Template Modeling score (TM-score) is also bounded between 0 and 1, with values above 0.5 generally indicating correct global topology, while scores below 0.17 suggest random similarity [33]. The Global Distance Test - Total Score (GDT-TS) ranges from 0 to 100, where higher percentages reflect greater structural overlap; values above 50 are usually considered structurally reliable in CASP evaluations [34]. Finally, the Predicted Aligned Error (PAE), introduced in AlphaFold2, outputs an expected error in Å for each residue pair, with lower values (typically below 5 Å) reflecting high confidence in relative positioning, while higher values signal uncertainty, especially across flexible domains [6].

For evaluating protein–protein interactions and docking quality, different specialized metrics are used. DockQ is a composite score that integrates interface RMSD (iRMSD), ligand RMSD (lRMSD), and the fraction of native contacts (Fnat) into a continuous scale between 0 and 1; values above 0.8 correspond to high-quality models, while scores below 0.23 are considered incorrect [35]. Fnat itself ranges from 0 to 1, where values above 0.5 suggest that at least half of the native intermolecular contacts are correctly predicted [36]. lRMSD, defined as the RMSD of the ligand after superimposing the receptor, is reported in Å; values below 2.5 Å generally indicate accurate docking poses, while values above 10 Å reflect incorrect placements [37]. iRMSD, which measures deviations only at the binding interface, is also expressed in Å; values below 2 Å correspond to high-quality predictions, while values above 4 Å suggest poor interface accuracy [37]. Together, these metrics cover both global orientation and fine-grained interaction fidelity in docking assessments.

### S3 Methods

#### S3.1 Environment Setup

A local setup was employed for downloading and constructing the unfolded protein structures. The local machine was equipped with an AMD Ryzen 7 5700U processor with Radeon Graphics (496 MB), 16 GB of RAM, and 477 GB of disk space, of which 438 GB was already occupied. For model training, a T4 GPU (15 GB) provided by Google Colab was utilized. A Pro subscription was employed to enable access to high-memory instances, allowing up to 50.1 GB of RAM.

#### S3.2 Preparing Proteins

As described in the main text, the model translates from the unfolded structure to the unique folded structure. Here, the methods used to obtain the ground truth folded structures and to construct the unfolded protein structures are detailed.

##### S3.2.1 Collection of Ground Truth Proteins

Protein structures were obtained from the RCSB Protein Data Bank (PDB) [112]. Due to limited local storage capacity (Sec. S3.1), only 40,000 proteins were downloaded. From this set, only pure peptide structures were retained, as PeptideBuilder [25], used to construct unfolded states, can process only pure peptides. Finally, as discussed in Sec. S1, structures containing hydrogen atoms are particularly important for model interpretability and accuracy. Consequently, the dataset was further restricted to Nuclear Magnetic Resonance (NMR) structures, which provide experimentally determined hydrogen positions. After filtering, a final dataset of 5,548 proteins was obtained, of which 20% (1,109) were reserved for testing, 10% (555) for validation, and the remaining 70% (3,884) for training.

The distribution of protein sequence lengths is shown in Fig. S1. Fig. S1.a depicts total sequence lengths, while Fig. S1.b illustrates chain-length distributions. Most proteins used for training have moderate lengths of approximately 100–200 residues, with some extending beyond 400, and a similar trend is observed in the chain-length distribution.

**Fig. S1:**
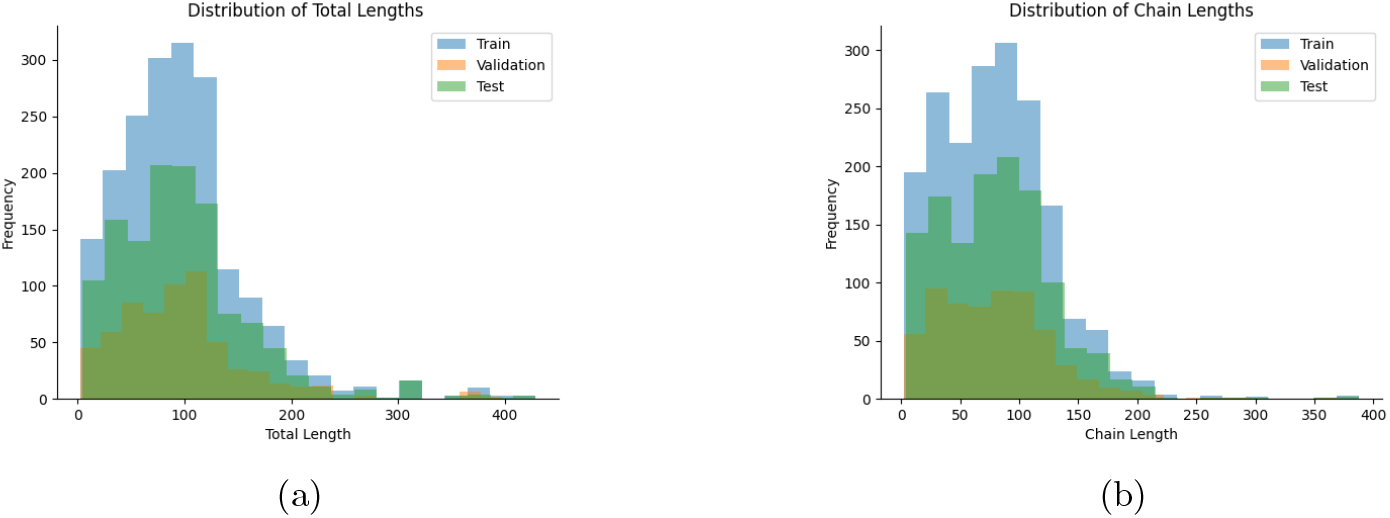
**a**. The total sequence length of the proteins that were used in training and testing. **b**. The total length of chains within each of those proteins. It should be mentioned that, ProteusFold is able to predict the structure of a protein without any explicit knowledge of chains.

##### S3.2.2 Construction of Unfolded Protein Structures

Unfolded conformations of each protein are generated using the PeptideBuilder [25] Python library. The amino acid sequence was extracted from the PDB file ^14^ and reconstructed with uniform backbone dihedral angles corresponding to an extended chain conformation:

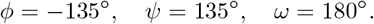

Here, *ϕ* is the torsion angle around the N–C_*α*_ bond, *ψ* is the angle around the C_*α*_–C bond, and *ω* is the angle around the peptide bond (C–N). These three dihedral angles fully define the local geometry of the polypeptide backbone, and setting them to extended values produces an idealized, unfolded chain. After construction, chain identifiers are reassigned to match the original PDB, TER records are inserted, and hydrogens are added using `PDBFixer` [26]. `PDBFixer` ensures chemically complete, simulation-ready structures by repairing missing atoms, replacing non-standard residues, and protonating the model. This process guarantees that each protein starts in a fully extended, unfolded state suitable for downstream modeling.

##### S3.2.3 Converting into Dataframes and Aligning

Following structure generation, both unfolded (ideal) and folded (first model) conformations are converted into node-level dataframes using BioPandas [113], with each row corresponding to an atom along with its residue identifiers and Cartesian coordinates. Because atom ordering and residue numbering^15^ are not consistent between the unfolded and folded structures, alignment is performed by matching residues and atom identities. Temporary residue identifiers are first assigned sequentially based on order (as the residue sequence is identical for both conformations), followed by consistent renumbering of residues across chains^16^. To maintain consistency, the atom ordering within each residue of the original folded protein is preserved and applied to the unfolded structure; therefore, residue and atom indices in the unfolded structure follow the folded structure’s convention^17^. The two dataframes are then merged into a single per-protein dataframe, ensuring that unfolded and folded coordinates are available side by side. All per-protein dataframes are subsequently concatenated into a unified dataset for downstream analysis.

Finally, arbitrary global rotations and translations between the folded and unfolded conformations are removed by applying the Kabsch algorithm [27] to optimally superimpose the folded coordinates onto the unfolded reference. This alignment ensures that all proteins are consistently oriented with respect to the unfolded backbone standard, placing both the start distribution and target folded distribution in a standardized and deterministic form.

#### S3.3 Tokenization Process

##### S3.3.1 Protein Structure Tokenization

###### Tokenization

A tokenization scheme is devised to standardize the representation of each residue while preserving physical constraints such as bond continuity. Let a protein consist of *N* residues, and let the *i*-th residue contain *n*_*i*_ *≤N*_max_ atoms. Let 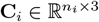 denote the absolute coordinates of the atoms in residue *i*. The representative structure token for residue *i* is defined as a matrix

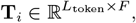

where *L*_token_ = *N*_max_ + 1 = 33 is the fixed token length and *F* = 3 corresponds to Cartesian coordinates (*x, y, z*). In the implementation, **T**_*i*_ is stored in `coord_tokens`, and an accompanying binary mask 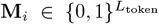 is stored in `coord_mask` to distinguish valid rows from padding (see Alg. 1).

Population of the token is performed as follows. The first *n*_*i*_ rows are filled with the residue coordinates and the remaining rows up to *N*_max_ are zero-padded:

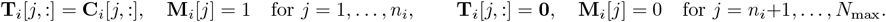

If a subsequent residue *i* + 1 exists, the synapse row (final row *L*_token_ = *N*_max_ + 1) is populated with the coordinates of the first atom of residue *i* + 1:

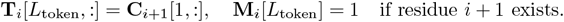

For normalization, each token is translated so that the first atom lies at the origin:

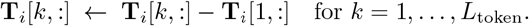

The mask ensures that downstream computations (e.g., attention or loss aggregation) consider only valid atomic or synapse rows; padded rows remain exactly zero after translation and therefore do not contribute spurious signal. This yields consistent residue-level tokens that capture local geometry and sequential connectivity along the chain, as illustrated in Fig. 1.b.

###### Detokenization

Let 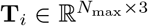 denote the token matrix for residue *i*, where **T**_*i*_[: *N*_*i*_,:] are the per-atom coordinates in the residue-local frame and **T**_*i*_[−1,:] is the synapse offset (the vector from residue *i* start to residue *i* + 1 start). Let **s**_*i*_ *∈* ℝ ^3^ denote the absolute start coordinate of residue *i*, stored in the array first atoms (with **s**_1_ = (0, 0, 0) when the protein origin is used).

Absolute start coordinates are recovered iteratively as

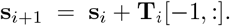

The absolute atomic coordinates of residue *i* are then obtained by translating the token coordinates by the recovered start:

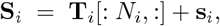

where *N*_*i*_ is the true atom count for residue *i*. After reconstruction, padding and synapse rows are removed, yielding a continuous set of absolute atomic coordinates that preserves residue connectivity and chain integrity.

##### S3.3.2 Atomic Properties Tokenization

For atomic embeddings, feature vectors 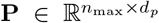 are derived from physico-chemical descriptors obtained using the Mendeleev library [28], where *d*_*p*_ denotes the descriptor dimension. Descriptors are extracted for protein-relevant elements (C, H, N, O, S, Se, D), with deuterium treated as hydrogen adjusted by isotope-specific values (e.g., atomic weight, neutron count, density). The resulting atomic tokens encompass properties such as atomic number, mass, electronegativity, radii, electron affinity, valency, density, polarizability, thermal conductivity, heat capacity, and neutron count.

All atomic features are first normalized using min–max scaling. The residues are then split into atomic tokens, each padded to a fixed length *L* _atom_ = 32, corresponding to the maximum number of atoms per residue in the dataset. After splitting, each token is shifted so that the nitrogen atom (N) occupies the origin. This centering provides a consistent reference frame, emphasizing relative positions and properties of other atoms (e.g., hydrogens retain meaningful values instead of being zeroed) and guiding the model to learn patterns of property variation relative to the backbone. The choice of nitrogen as the reference is user-defined; alternative atoms or normalization schemes could be adopted in future implementations depending on the modeling objective. The resulting tokens therefore provide a chemically meaningful and positionally standardized representation suitable for downstream structural learning.

#### S3.4 Encoder architecture

Both the structural and atomic encoders are implemented as compact multilayer perceptron (MLP) autoencoders that map variable-length, per-residue inputs to fixed-size latent embeddings. The presentation below gives a concise mathematical description, and explicit implementation details sufficient for exact reproduction.

##### Notation

Let a single residue input be *X ∈* ℝ^*L×F*^, where *L* is the maximum number of atom tokens per residue and *F* is the number of features per atom. Denote by 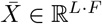 the row-major flattened vector of *X*. The encoder implements a mapping *f*_enc_: ℝ^*L·F*^ *→*ℝ^*d*^ producing a latent vector 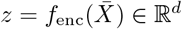. The decoder implements the inverse map *f*_dec_: ℝ^*d*^ *→*ℝ^*L·F*^ with reconstructed 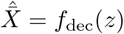 and 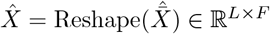.

##### Formal layer specification

The encoder is realized as a sequence of affine transforms and ReLU nonlinearities:

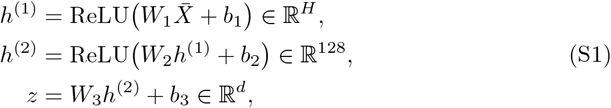

where *W*_1_ *∈*ℝ^*H×*(*L·F*)^, *W*_2_ *∈*ℝ^128*×H*^, *W*_3_ *∈*ℝ^*d×*128^ and *H* denotes the first hidden width. The decoder mirrors this topology:

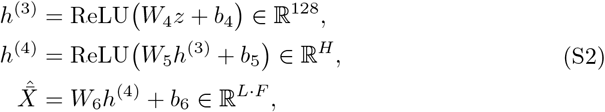

with 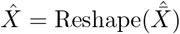.

##### Masked reconstruction loss

To accommodate variable atom counts per residue, let *M ∈ {* 0, 1 *}*^*B×L*^ be a binary mask over atoms in a batch of size *B* (1 = valid). A masked mean squared error was used:

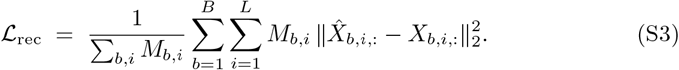

This avoids penalizing padded positions and yields stable gradients for variable-sized residues.

##### Implementation details

All encoder models were implemented in PyTorch [114] using a unified class named “AE,” through which a simple yet flexible MLP-based autoencoder is provided for both residue-level structural tokens and atom-level property vectors. Inputs are represented as batch tensors **X** *∈* ℝ^*B×L×F*^, where (B) denotes the batch size, (L) the maximum token length, and (F) the feature dimensionality. For the structural encoder, (F=3) (Cartesian coordinates) and *L*_struct_ = 33 are used, while for the atomic encoder, (F=13) and *L*_atom_ = 32 are specified. Prior to encoding, inputs are flattened to shape (*B, L ·F*) using nn.Flatten(). The encoder is constructed with two hidden layers of widths (256) and (128), followed by a latent layer of dimension (d=128), resulting in the mapping *L· F →* 256 *→* 128 *→*128. The decoder is designed to mirror this structure (128 *→* 128 *→* 256 *→ L · F*) and employs linear activation in the final layer for regression to the original feature space, while ReLU activations are applied in all intermediate layers. In the forward pass, both the reconstructed tensor 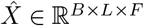 and the latent representation *z ∈* R^*B×d*^ are returned. To handle variable-length inputs, sequences are padded to length (L), and reconstruction loss is computed with the aid of masks (M), ensuring that positions with (M=0) do not contribute to the masked MSE loss.

##### Encoder Training Details

The structural autoencoder, with input feature dimension *F* = 3 (Cartesian coordinates), token length *L* = 33 (*N*_max_ + 1 = 33), and latent dimension *d* = 128, is trained using the Adam optimizer with an initial learning rate of 1 × 10^−3^. A StepLR scheduler is applied to gradually reduce the learning rate by a factor of 0.8 every 20 epochs, promoting stable convergence. Training is performed for 300 epochs with a batch size of *B* = 512, ensuring efficient optimization and consistent reconstruction across the dataset.

For the atomic property autoencoder, the input feature dimension is *F* = 13 and the token length is *L* = 32 (*N*_max_ = 32). All other hyperparameters remain the same, except the number of epochs, which is set to 5 due to the limited number of 22 amino acids present in the proteins.

#### S3.5 Preparing Sequence

All the residue encoded tensor were converted into fixed-length, protein-level sequences to enable batch processing. Each residue embedding was represented as *x*_*i*_ *∈* ℝ^*d*^, where *d* denotes the feature dimension. For a protein *p* containing *n*_*p*_ residues, the sequence is written as

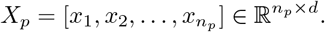

To standardize sequence lengths, each *X*_*p*_ was either truncated or zero-padded to a fixed maximum length *L*_max_ = 400. A binary mask 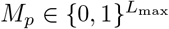 was constructed in parallel, where valid residues were marked with 1 and padded positions with 0.

Finally, the processed dataset was stacked into tensors of shape 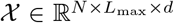 and 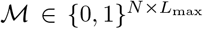, where *N* denotes the number of proteins. The use of zero padding ensured that padded positions did not contribute to model computations. A value of *L*_max_ = 400 was adopted, as the majority of proteins in the dataset fell within this range (Fig. S1), although longer limits may be employed in future work.

#### S3.6 Translation

##### S3.6.1 Intuition of the translation process

The translation process used a Transformer named as “DualTransformerWithAttn” that, with the guidance of atom-level information, translates the unfolded residue-level structural tokens into the folded representations in two conceptual stages. First, the encoder ingests a sequence of atomic-property embeddings (**X**_src_) and builds a *memory* that summarizes local and long-range atomic context across the whole sequence. Second, the decoder consumes unfolded residue token embeddings (**X**_tgt in_) and, for each residue, (i) uses self-attention to model target-side dependencies and (ii) uses cross-attention to query the encoder memory so that residue predictions are grounded in atom-level context. Attention maps emitted by the model reveal which source positions the decoder queries for each predicted target token.

##### S3.6.2 Mathematical formulation

Notation: *L*_seq_ = 400 (max sequence length), *D* = 128 (embedding dimension), batch size *B*. Positional embeddings 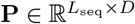 are added to inputs.

Encoder input (positionalized):

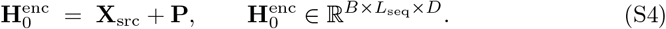

A single encoder layer *l* applies multi-head self-attention (MHSA) and a feed-forward network (FFN) with residual connections and layer normalization:

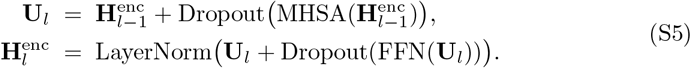

The final encoder output 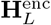 is called the *memory*.

Decoder input (positionalized):

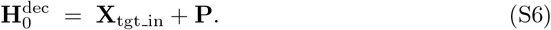

A single decoder layer *l* applies decoder self-attention, cross-attention over encoder memory, and an FFN:

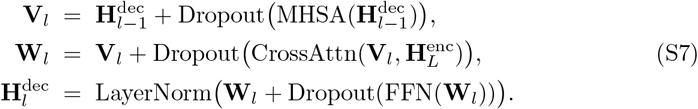

Final projection to predicted structural token embeddings:

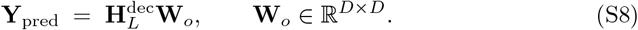

Masking: boolean masks **M**_src_,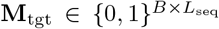 indicate valid positions; for PyTorch’s ‘MultiheadAttention’ the key-padding mask is the logical inverse (True = padded). Attention maps returned per layer have shapes (*B, L*_seq_, *L*_seq_) for self-attention and (*B, L*_seq_, *L*_mem_) for cross-attention.

##### S3.6.3 Implementation details

The model was implemented in PyTorch, and its components are specified here in sufficient detail to enable exact reproduction. Input sequences were first embedded using a positional embedding layer (`nn.Embedding`) with a maximum sequence length of *L*_seq_ = 400 and embedding dimension *D* = 128.

The encoder consisted of two stacked encoder blocks named “`EncoderLayerWithAttn`”. Each block employed a multi-head self-attention mechanism with four heads (`nn.MultiheadAttention`, embed dim = 128, `batch first=True`), followed by residual connections, layer normalization (`nn.LayerNorm(128, eps=1e-5)`), and a feed-forward network of dimension 128 *→* 512 *→* 128 with ReLU activation and dropout (*p* = 0.1).

The decoder mirrored this structure and was composed of two decoder blocks named “`DecoderLayerWithAttn`”. Each block applied self-attention over the decoder input, cross-attention over the encoder memory, and a feed-forward network of identical form to the encoder (128 *→* 512 *→* 128). All sublayers were followed by residual connections, dropout (*p* = 0.1), and layer normalization (`nn.LayerNorm(128)`).

Finally, a linear projection layer (`nn.Linear(128, 128)`) mapped the decoder outputs to the target embedding space. This configuration—positional embedding of size 400 × 128, two encoder layers, two decoder layers, four attention heads, feed-forward dimension of 512, and dropout of 0.1—was used consistently across all experiments reported in this study.

###### Training and Hyperparameter Configuration

The DualTransformerWithAttn model was instantiated with 128-dimensional embeddings, 4 attention heads, 2 encoder and 2 decoder layers, a feed-forward hidden dimension of 512, dropout rate of 0.1, and maximum sequence length *L*_max_ = 400:

Training employed the Adam optimizer with an initial learning rate of 1 × 10^−4^. Learning rate adaptation was performed using a `ReduceLROnPlateau` scheduler, which reduced the learning rate by a factor of 0.8 after 20 epochs of stagnating validation loss, with a lower bound of 1 × 10^−7^ to prevent vanishing updates.

The model was trained with a batch size of *B* = 32, using standard splits of approximately 70%/10%/20% for training, validation, and testing, respectively. Reproducibility was ensured by setting a fixed random seed (42) via `torch.manual seed(seed)`.

The loss function was a masked mean squared error:

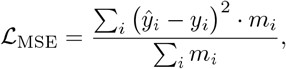

where *m*_*i*_ indicates valid (non-padded) positions. Dropout of 0.1 was applied for regularization. The model was trained for 570 epochs, which allowed sufficient convergence while ensuring stability in both training and validation metrics.

##### S3.6.4 Reconstructing the Protein Structure

The predicted embeddings are passed through the structural decoder to obtain padded structural tokens. These tokens are then detokenized into Cartesian coordinates, after which padding rows and the final row (Synapse) of each token are removed. The result is the reconstructed protein structure.

**Fig. S2:**
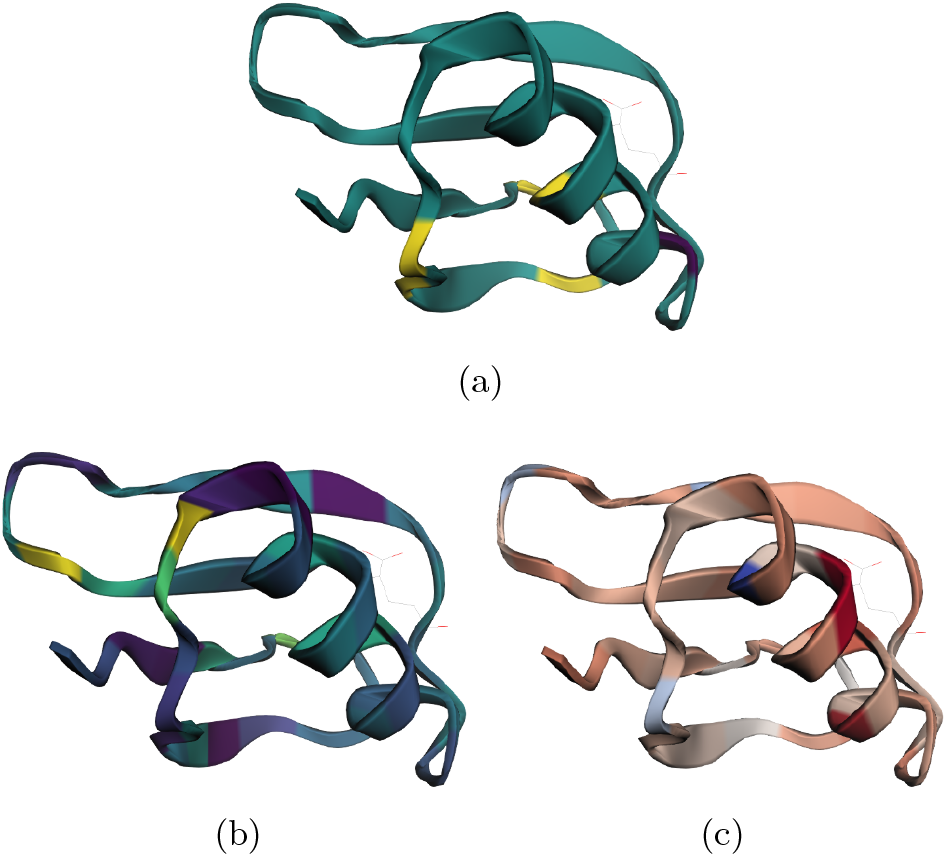
**a**. Interactive visualization representation of attention-given for the protein 1GB1. **b**. Interactive visualization representation of rollout-recieved for the protein 1GB1. **b**. Interactive visualization representation of gradient-based *gradient × input* for the protein 1GB1.

### S4 Explanation Collection

#### S4.1 Attention Collection

To quantify the role of each residue in the model’s attention mechanism, three metrics were extracted namely, *attention received, attention given*, and *rollout received*. These were derived exclusively from the encoder self-attention maps, as the encoder directly processes the residue-level atomic property embeddings (Sec. S3.3.2), making its attention patterns biologically interpretable. Decoder cross-attention weights were also collected during training runs but are not reported here, since they represent alignment between unfolded latent sequence and target folded space rather than intra-residue relations.

##### Attention preprocessing

For each protein input, encoder self-attention maps are extracted from all layers, averaged across attention heads, and subsequently aggregated across layers. Invalid (padded) positions are masked to ensure normalization across only valid residues. In all attention matrices, rows correspond to query residues giving attention, and columns correspond to key residues receiving attention.

- **Attention-given:** For a query residue *i*, this is computed as the row-sum of the normalized attention matrix *A*, representing the total attention distributed by residue *i* to other residues. A visual representation of the interactive 1GB1 protein structure, with attention-given shown as a color mapping, is presented in Fig. S2.a.
- **Attention-received:** For a key residue *i*, this is computed as the column-sum of *A*, representing the total attention directed toward residue *i* from all other residues. A visual representation of the interactive 1GB1 protein structure, with attention-received shown as a color mapping, is presented in Fig. 4.a.
- **Rollout-received:**To capture multi-layer information flow, the attention rollout procedure is applied, where each layer’s attention matrix is augmented with an identity matrix, row-normalized, and multiplied sequentially. The resulting joint matrix encodes effective connectivity between residues across layers. The *rollout-received* score for residue *i* is defined as the sum of the *i*-th column of this matrix, representing the cumulative information reaching that residue. A visual representation of the interactive 1GB1 protein structure, with rollout-received values shown as a color map, is presented in Fig. S2.c.

All residue-level scores were stored in a structured dataframe indexed by protein and residue identifiers, enabling systematic aggregation, downstream statistical analysis, and biological visualization.

##### S4.1.1 Understanding the Attentions

As mentioned in the main text, it is most advisable to rely on the primary attention matrix for interpretation. For added nuance, however, the three derived values described above can also be useful, particularly in cases where more detailed analysis is required. While this study focused primarily on residues that stand out most clearly, future researchers may obtain a more refined perspective by incorporating these derived measures.

The three values provide complementary insights into residue-level attention. “attention-received” highlights residues most strongly influenced by others, reflecting their relative importance within the folding context. “rollout-received” offers a normalized view, enabling comparisons across residues regardless of overall scale. In contrast, “attention-given” identifies residues that exert the greatest influence on others, potentially revealing those that drive structural organization. Together, these measures enrich the interpretation of the attention mechanism by balancing the identification of key residues with insights into their directional influence within the protein.

#### S4.2 Residue-level attribution via gradient×input

To interpret residue-level importance, a *gradient × input* approach is applied, which is categorized within the family of gradient-based explanation techniques [45, 46]. Let *x ∈* ℝ^*d*^ denote the residue-level input features and let *y* = *f* (*x*) be the model output. The attribution for feature *x*_*i*_ is defined in Eq. S9, quantifying how a small perturbation of *x*_*i*_ influences the prediction *y*, scaled by the magnitude of the input itself.

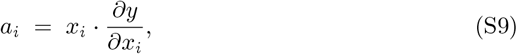

However, in this case *y* is treated as a latent representation of protein structure, for which increases or decreases in individual coordinates are not directly interpretable. To address this, attributions are computed with respect to a supervised training objective, namely the mean squared error (MSE) between the prediction *y*_pred_ = *f* (*x*) and a reference representation *y*_ref_, as defined in Eq. S10.

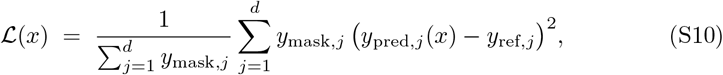

Here, *y*_mask_ *∈* {0, 1 }^*d*^ is used as a binary mask to select only valid (non-padded) positions. This ensures that only meaningful residues contribute to the gradient signal, and normalization by ∑_*j*_ *y*_mask,*j*_ accounts for differences in sequence length.

The gradient of the loss with respect to the input is shown in Eq. S11.

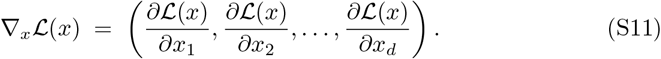

Finally, the gradient × input attribution with respect to the loss is defined in Eq. S12.

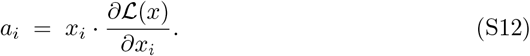

Equation S12 extends the basic definition in Eq. S9 by grounding the attributions in the training objective (Eq. S10). Here, *a*_*i*_ < 0 is interpreted as indicating that the input feature *x*_*i*_ helps move the prediction closer to the reference structure, whereas *a*_*i*_ > 0 suggests that the feature pushes the prediction farther away.

#### S4.3 Atomic-level attribution via gradient×input

Model predictions are attributed to atomic properties using a two-stage gradient pipeline that follows the gradient × input principle (Eq. S9) and the masked loss (Eq. S10).

In the first stage, per-residue gradients are collected from the trained sequence model. For each residue *r*, the upstream gradient is denoted 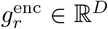, obtained by backpropagating the scalar loss through the residue embeddings. These gradients are stored for all residues in a protein.

In the second stage, the atomic feature tensor **X** *∈* ℝ^*R×L×P*^, where *R* is the number of residues, *L* is the number of atoms per residue, and *P* is the number of atomic properties, is passed through the atomic encoder to produce residue embeddings **E** = enc_atom_(**X**). The stored residue gradients 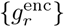 are used as upstream signals in a vector–Jacobian product (VJP) [65, 66] to compute gradients with respect to the atomic inputs:

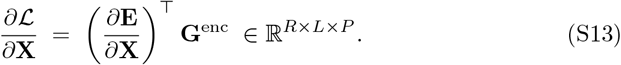

Atomic attributions are then computed using the gradient×input rule:

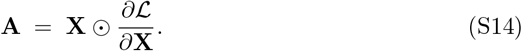

Here, **A**_*r,ℓ,p*_ represents the signed attribution for property *p* of atom *ℓ* in residue *r*. Only valid atoms are allowed to contribute, enforced by multiplying each atom by its mask *M*_*r,ℓ*_ *∈ {* 0, 1 *}*, i.e., **Ã** _*r,ℓ,p*_ = *M*_*r,ℓ*_**A**_*r,ℓ,p*_.

These atomic attributions are aggregated to produce per-residue or per-protein summaries. For example, the sum or absolute sum over atoms in a residue is computed as 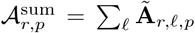 or 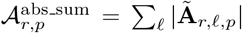. Mean aggregation is performed by dividing by the number of valid atoms.

## S5 Location of Interest Identification

To quantify the relationship between attention signals and local structural irregularities, *locations of interest* (termed *anchors* in the code) were identified for each protein sequence. A location of interest is defined as a short, contiguous interval of residues centered on a pivot residue (the pivot is a high-attention residue). Two complementary strategies were supported: (i) user-specified ranges (when available) and (ii) automatic detection.

Formally, let a protein *p* have residues indexed by integers and let *S*_*p*_ = *{*1, …, *n*_*p*_*}* denote its residue index set. A location of interest for *p* is specified as an interval

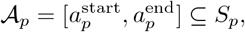

where 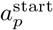 and 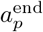 are inclusive residue indices. If a user-provided range is available, that interval is used directly. Otherwise the pivot residue *π*_*p*_ is determined automatically as described below and a symmetric window of radius *w* is taken:

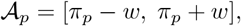

with *w* a hyperparameter (default *w* = 2).

### Automatic pivot selection

When an anchor is not provided, the pivot *π*_*p*_ is derived from the rollout attention statistics aggregated across samples of the same protein. Let the dataset contain *S* independent samples (e.g., different masking / batching instances) for protein *p*. For each sample *s* an index *r*_*p,s*_ corresponding to the residue with maximum rollout score is identified (ties are broken deterministically). The pivot is then chosen as the most frequently occurring top residue across samples:

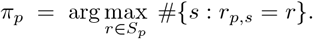

If no clear counts are available (e.g., all *r*_*p,s*_ are missing), the median residue index median(*{* Residue ID *}*) is used as a fallback pivot.

### Per-sample top-residue membership and summary statistics

For each sample of protein *p*, the residue with highest encoder attention (or highest rollout score) is identified; let these per-sample top indices be denoted 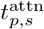 and 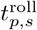, respectively. A sample is counted as *aligned* if the top residue falls inside the anchor interval: *t∈𝒜* _*p*_. Over the set of valid samples 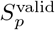 (samples for which top-residue indices are available), the alignment percentages are reported as

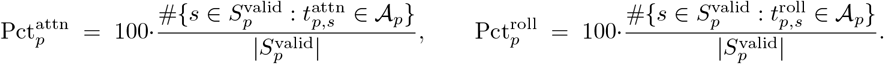

These percentages are reported per protein for both direct attention and rollout attention.

### Practical details

In implementation (`function_compute_anchor_top_percentages`), input attention tables are validated to contain required fields (including `dataset_idx, Residue ID, attn_received`, and rollout received); non-numeric residue IDs are coerced to integer-like values and missing entries are handled by skipping affected samples. When user-specified anchors are supplied (a mapping {`protein id: (start,end)` }), they take precedence over automatically derived anchors. Proteins with fewer than a user-defined minimum number of samples (parameter `min samples threshold`, default = 1) can be flagged or excluded from aggregated summaries. All per-protein results (including *n*_samples_, number of valid samples, anchor bounds, anchor derivation method, raw counts and percentages for attention and rollout alignment) are written to a CSV file for downstream analysis.

The procedure as implemented thus provides a simple, reproducible mechanism to (i) define locations of interest either from prior knowledge or from attention-derived pivots, (ii) evaluate whether per-sample high-attention residues fall within those locations, and (iii) summarize alignment rates across the dataset. The code for this procedure (including tie-breaking rules, NaN handling, and CSV export) is provided in the repository and was used to generate the statistics reported in the manuscript.

### S6 Intuitive Interpretation of Gradient-Based Explanations under MSE Loss

The detailed procedures for computing gradient × input at the residue and atomic levels are provided in Sec. S4.2 and Sec. S4.3. Here, a generalized explanation is provided to give an intuitive understanding of how these values should be interpreted when a regression model is trained with mean squared error (MSE).

The MSE loss is defined as:

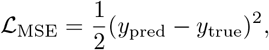

which measures the deviation of the prediction from the true value.

The gradient with respect to the prediction is obtained as:

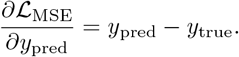

By applying the chain rule, the gradient with respect to each input feature *x*_*i*_ can be expressed as:

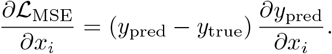

The gradient × input method then multiplies this sensitivity by the feature value itself:

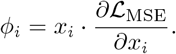

### Intuitive interpretation

This process can be understood by imagining each feature *x*_*i*_ as a dial on a control panel: - A large |*ϕ*_*i*_| indicates that the dial has a strong effect on the prediction error. - A positive value (*ϕ*_*i*_ > 0) indicates that increasing the feature would increase the loss, thereby moving the prediction further away from the true value. - A negative value (*ϕ*_*i*_ < 0) indicates that increasing the feature would reduce the loss, thereby moving the prediction closer to the true value.

Thus, positive and negative values should not be interpreted as “good” or “bad” features; instead, they indicate the direction of influence that each feature exerts on the prediction relative to the ground truth.

### S7 Further Discussion on Atomic Properties

Fig. S3.a shows a pie chart indicating that atoms in all proteins are dominated by hydrogen and carbon, with oxygen and nitrogen comprising a modest portion. Although sulfur atoms are few in number, as will be shown, their influence on the protein is very significant. As illustrated in Fig. S3.b, most properties—except for specific heat (dominated by hydrogen) and thermal conductivity (dominated by carbon)—are largely influenced by sulfur. Fig. S4.a shows that sulfur’s electron affinity exhibits moderate variation, with a notably high spread in dipole polarizability. Sulfur also exerts considerable influence over electronegativity, covalent radius, and atomic radius. While sulfur’s electronegativity (2.58 on the Pauling scale) is significantly lower than oxygen’s (3.44), its larger covalent and atomic radii combined with high polarizability enhance its ability to form unique, stabilizing interactions within proteins. Refer to Tables S1 and S2 for the numerical values.

**Fig. S3:**
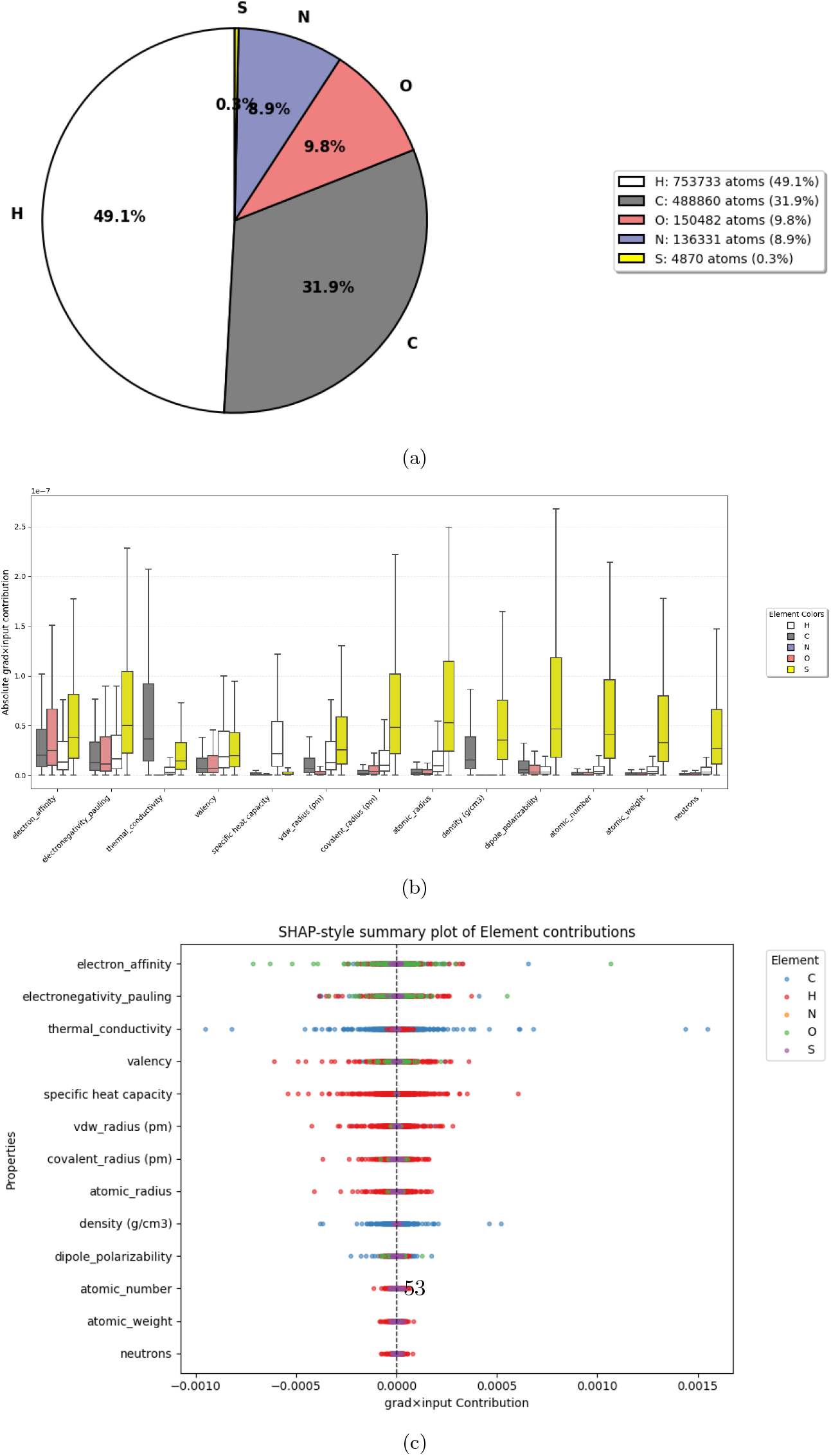
**a**. Pie Chart showing the comparison of the count of each element in the proteins. **b**. Box plot of the absolute *gradient × input* values for all atomic properties, grouped by element. **b**. Shap like visualization of *gradient × input* importance of all Atomic properties for each element.

**Table S3:** List of the 22 proteinogenic amino acids with their names, three-letter, and one-letter codes.

| Amino Acid | Three-letter Code | One-letter Code |
| --- | --- | --- |
| Alanine | Ala | A |
| Arginine | Arg | R |
| Asparagine | Asn | N |
| Aspartic acid | Asp | D |
| Cysteine | Cys | C |
| Glutamic acid | Glu | E |
| Glutamine | Gln | Q |
| Glycine | Gly | G |
| Histidine | His | H |
| Isoleucine | Ile | I |
| Leucine | Leu | L |
| Lysine | Lys | K |
| Methionine | Met | M |
| Phenylalanine | Phe | F |
| Proline | Pro | P |
| Serine | Ser | S |
| Threonine | Thr | T |
| Tryptophan | Trp | W |
| Tyrosine | Tyr | Y |
| Valine | Val | V |
| Selenocysteine | Sec | U |
| Pyrrolysine | Pyl | O |

Several studies support that sulfur’s high polarizability and electron affinity enable it to form strong noncovalent interactions—such as with oxygen, nitrogen, and aromatic rings—that stabilize protein folding [84–86]. Methionine and cysteine residues contribute through disulfide bonds and chalcogen bonding, which are critical for maintaining tertiary protein structure stability [84, 87].

Fig. S3.c shows a SHAP-like summary of the *gradient × input* values for the elements, highlighting hydrogen’s dominance in these properties. This confirms that, collectively, atomic properties are dominated by hydrogen and carbon due to their abundance, whereas sulfur, despite its scarcity, plays a disproportionately significant role.

Fig. S4.b presents mean property values, which do not capture the full extent of their influence. Proper analysis requires examining individual atoms for accurate interpretation. Notably, the electron affinity of most carbon atoms tends to have a destabilizing (positive) effect, while sulfur exhibits a stabilizing (negative) effect on protein folding, consistent with prior discussions. Similar patterns are observed for thermal conductivity: carbon’s influence is high but destabilizing, inducing local fluctuations in the fold, as previously reported [80]. Sulfur, on the other hand, shows a slight stabilizing impact. In contrast, sulfur generally exerts a destabilizing effect on protein folding in most other properties. Further studies need to be conducted to verify the claims.

**Fig. S4:**
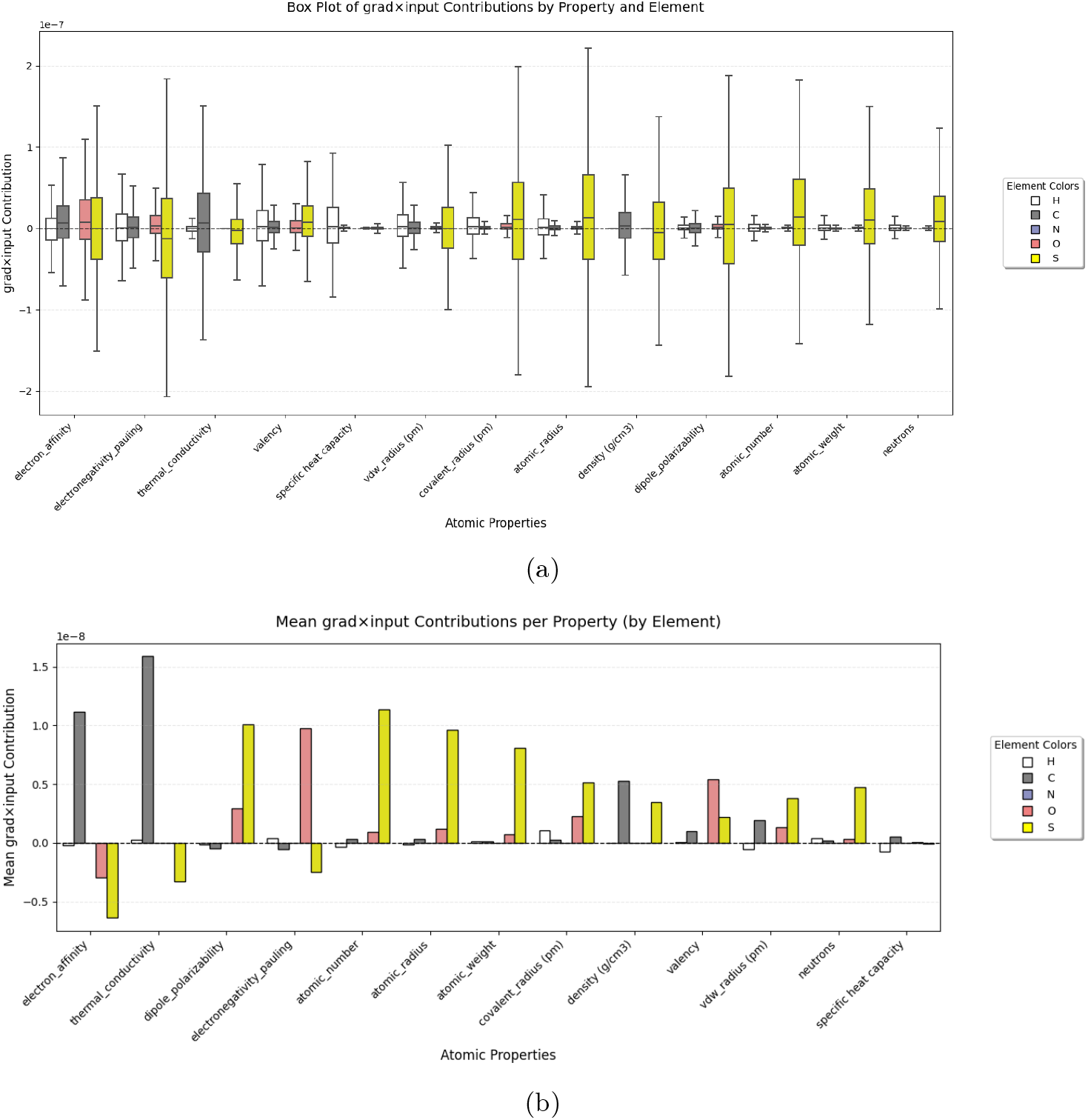
**b**. Box plot of the true *gradient × input* values for all atomic properties, grouped by element. **b**. Histograms showing the meaning of each properties. It should be noted that these are the mean so it doesn’t represent the entire picture of their influence.

**Fig. S5:**
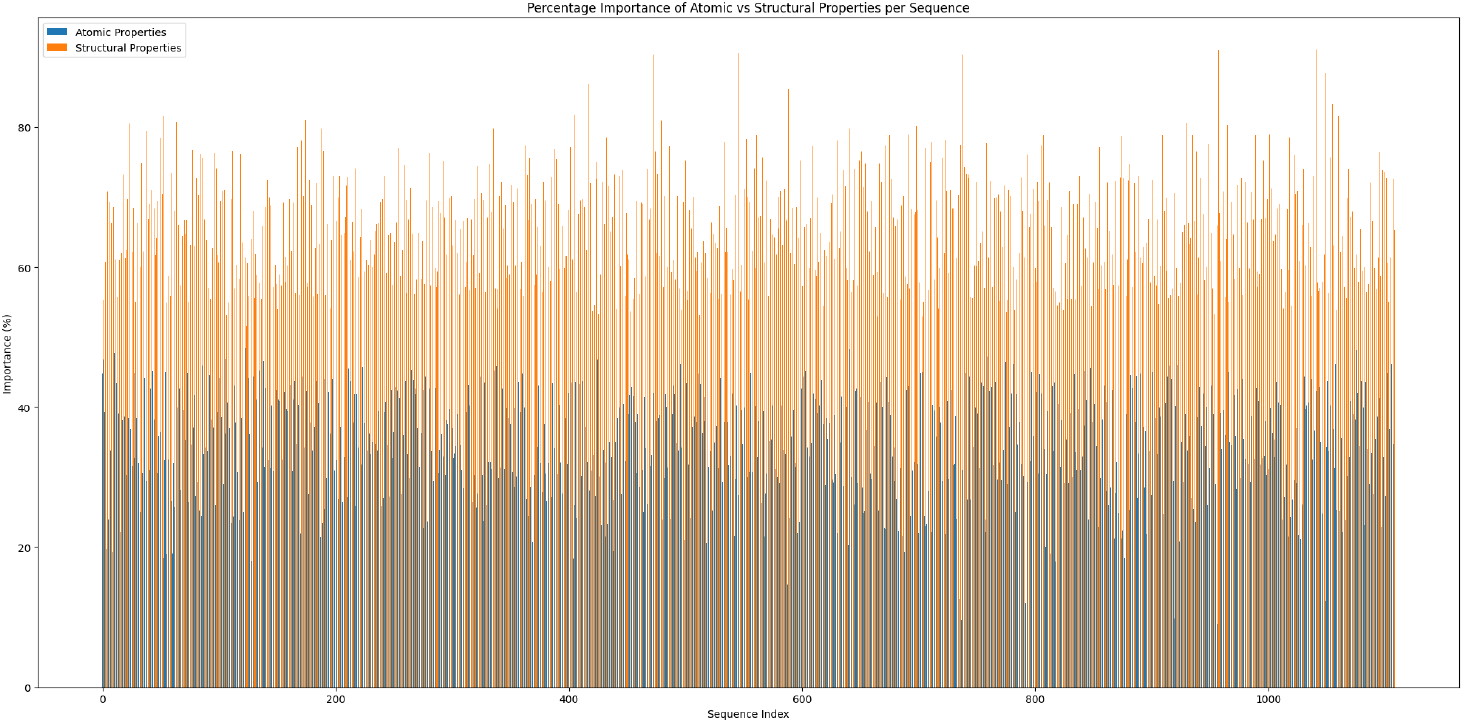
Comparison between the *gradient × input* between Unfolded embedding sequence and Atomic properties embedding sequence.

### S8 Brief discussion on the two input embeddings

Fig. S5 compares the *gradient × input* attributions of the Unfolded Structure Embedding Sequence and the Atomic Properties Embedding Sequence. Unlike earlier *gradient × input* analyses, which were computed on the MSE loss, this calculation is based on the prediction itself and therefore should not be interpreted in terms of positive or negative effects on prediction quality. Instead, it provides a rough estimate of the relative proportional influence of the two embeddings. The contributions are 67.69% for the unfolded structure and 32.31% for the atomic properties, indicating that structural tokens contribute more heavily, as they directly encode the initial residue state. Nonetheless, atomic property embeddings remain substantially informative, providing complementary signals that enrich residue characterization.

### S9 Approximate Cost Estimation

To contextualize the computational efficiency of the method in this study (*ProteusFold*), its estimated costs were compared against three major structure prediction frameworks: AlphaFold2, RosettaFold, and OpenFold. Cost estimation was based on two components: (i) electricity usage, derived from GPU/TPU power ratings and training duration, and (ii) purchase or subscription costs. Electricity costs were calculated assuming an average U.S. commercial electricity price of $0.1622/kWh. [115] It should be noted that these cost estimates correspond to the time of publication and are approximate, so the actual values may differ. It is also disclosed that a total of 400 units were purchased from Google Colab, amounting to approximately $44. Most of these units were consumed in failed attempts, with a significant portion spent on obtaining results from AlphaFold2. However, for the purpose of comparison, only the estimated training costs are reported here.

The configurations reported in the original papers were used to approximate the costs. AlphaFold2 was executed on 128 TPU v3 devices for a total of 11 days (264 hours) [7]. RosettaFold computations were carried out using 8 NVIDIA V100 GPUs (32 GB each) over 28 days (672 hours) [10]. OpenFold was run on 128 NVIDIA A100 GPUs for 8 days (192 hours) [13], whereas ProteusFold was evaluated on a single NVIDIA T4 GPU, with training lasting approximately 3 hours.

The energy consumption of T4 as shown in their offical site is 70W [116], A100 was seen to be 400W[117], V100 was seen to be 300 [118] and TPUv3 has 262W [119] using these information the energy cost was calculated. Table S4 shows the energy consumption of each model. AlphaFold2 consumed 8,853.5 kWh, RosettaFold 1,612.8 kWh, OpenFold 9,830.4 kWh, while ProteusFold required only 0.21 kWh.

The purchase cost of GPUs was obtained from public sources: the NVIDIA A100 at $10,000.00 [120] and the V100 at $1,570.00. For AlphaFold2, training was originally performed on TPU v3 pods. As public information on the hardware purchase cost of TPU v3 nodes is unavailable, the Google Cloud TPU v3 subscription pricing of $2.20 per hour per device [121] was used to approximate the compute cost. For ProteusFold, experiments were conducted using Google Colab Pro, which is priced at $11 for 100 compute units. On average, 1.2 units corresponded to 1 GPU-hour for a T4, giving an effective rate of

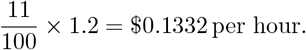

This rate was multiplied by the runtime hours to obtain the final cost reported in Table S5. Table S5 combines hardware/subscription and electricity costs. The total costs incurred were $75,778.44 for AlphaFold2, $12,821.60 for RosettaFold, $1,281,594.49 for OpenFold, and $0.42 for ProteusFold.

These results demonstrate the stark contrast between large-scale structure prediction models and the lightweight approach presented here. While thousands of dollars in resources and megawatt-scale energy usage were required by AlphaFold2 and Open-Fold, comparable interpretability results were achieved by ProteusFold with negligible energy consumption and total costs below one dollar, highlighting its accessibility and sustainability.

**Table S4:** Energy consumption and electricity costs for each model. Electricity price assumed at $0.1622 per kWh (USA). AlphaFold2 consumed 8,853.5 kWh ($1,436.04), RosettaFold 1,612.8 kWh ($261.60), OpenFold 9,830.4 kWh ($1,594.49), and ProteusFold only 0.21 kWh ($0.034).

| Model | GPUs & Type | Hours | Power (W) | Energy (kWh) | Electricity Cost (\$) |
| --- | --- | --- | --- | --- | --- |
| AlphaFold2 | 128 TPUv3 | 11 × 24 = **264** | 262 | 262 × 128 × 264 / 1000 = **8,853.50** | 8,853.50 × 0.1622 = **1,436.04** |
| RosettaFold | 8 V100 (32GB) | 28 × 24 = **672** | 300 | 300 × 8 × 672 / 1000 = **1,612.80** | 1,612.80 × 0.1622 = **261.60** |
| OpenFold | 128 A100 | 8 × 24 = **192** | 400 | 400 × 128 × 192 / 1000 = **9,830.40** | 9,830.40 × 0.1622 = **1,594.49** |
| ProteusFold | 1 T4 | 3 | 70 | 70 × 3 × 1 / 1000 = **0.21** | 0.21 × 0.1622 = **0.034062** |

**Table S5:**
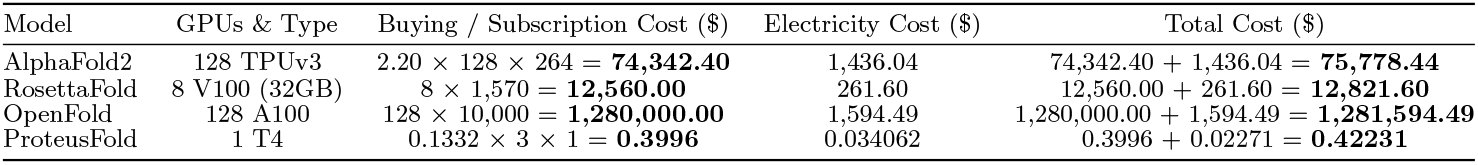
Total cost including compute subscription/buying and electricity. AlphaFold2 totaled $75,778.44, RosettaFold $12,821.60, OpenFold $1,281,594.49, while ProteusFold required only $0.42. Electricity cost column is imported from Table S4.

## Footnotes

1 Coordinates of atoms N, C*α*, C, O for each residue.

2 Each recycling iteration re-applies the network (with tied weights) to refine the structure by feeding previous predictions back into the model.

3 Protein language models are transformer-based architectures with self-attention layers, trained on millions to billions of protein sequences to capture contextual amino acid representations, analogous to words in natural language.

4 As noted earlier, AlphaFold2 addressed this issue with 200 relaxation cycles per prediction.

5 Here, *ϕ* is the torsion angle around the N–C*α* bond, *ψ* around the C*α*–C bond, and *ω* around the peptide bond (C–N).

6 Chain identifiers do not play any role in the model. They are added for convenience of understanding.

7 The figure shows the folded structure but the method is applied on both folded and unfolded structures.

8 The batch implementation of AlphaFold provides only PAE values. It should also be noted that this is the only version supporting batch processing; implementation of other models require per-sequence processing, which is computationally infeasible for large-scale evaluation.

9 The term SET domain refers to a conserved protein domain originally identified in three *Drosophila* proteins: *Su(var)3-9, Enhancer of zeste*, and *Trithorax*, from which the acronym “SET” is derived

10 IgG-Fc is the tail region of an IgG antibody that mediates effector functions, such as binding to Fc receptors and activating complement, while the Fab region binds antigens.

11 The rate-limiting transition state in protein folding is the high-energy intermediate conformation that the protein must cross, representing the slowest step that determines the overall folding rate.

12 H23C indicates substitution of histidine (H) at position 23 with cysteine (C).

13 Complement component 3, often simply called C3, is a protein of the immune system that is found primarily in the blood [88].

14 Alternatively, the sequence can be provided directly by the user.

15 Some protein residues have different residue IDs because they may belong to a fragment of the protein rather than the full sequence.

16 When interpreting the residues one should be aware of the initial numbering of the residues.

17 This convention is a design choice; the convention of unfolded structures can also be adopted if deemed suitable.

